# A mutational gradient drives somatic mutation accumulation in mitochondrial DNA and influences germline polymorphisms and genome composition

**DOI:** 10.1101/2021.06.18.448725

**Authors:** Monica Sanchez-Contreras, Mariya T. Sweetwyne, Brendan F. Kohrn, Kristine A. Tsantilas, Michael J Hipp, Elizabeth K. Schmidt, Jeanne Fredrickson, Jeremy A. Whitson, Matthew D. Campbell, Peter S. Rabinovitch, David J. Marcinek, Scott R. Kennedy

## Abstract

**Background:** Mutations in the mitochondrial genome (mtDNA) can cause devastating maternally inherited diseases, while the accumulation of somatic mtDNA mutations is linked to common diseases of aging. Although mtDNA mutations impact human health, the process(es) that give rise to these mutations are unclear and are under considerable debate. We analyzed the distribution of naturally occurring somatic mutations across the mouse and human mtDNA obtained by Duplex Sequencing to provide clues to the mechanism by which *de novo* mutations arise as well as how the genome is replicated.

**Results:** We observe two distinct mutational gradients in G→A and T→C transitions, but not their complements, that are delimited by the light-strand origin and the control region (CR). The gradients increase with age and are lost in the absence of DNA polymerase γ proofreading activity. A nearly identical pattern is present in human mtDNA somatic mutations. The distribution of mtDNA single nucleotide polymorphisms (SNPs) in the human population and genome base composition across >3,000 vertebrate species mirror this gradient pattern, pointing to evolutionary conservation of this phenomenon. Lastly, high-resolution analysis of the mtDNA control region highlights mutational ‘hotspots’ and ‘cold-spots’ that strongly align with important regulatory regions.

**Conclusions:** Collectively, these patterns support an asymmetric strand-displacement mechanism with key regulatory structures in the CR and argue against alternative replication models. The mutational gradient is a fundamental consequence of mtDNA replication that drives somatic mutation accumulation and influences inherited polymorphisms and, over evolutionary timescales, genome composition.

## Introduction

Owing to their evolutionary origin, mitochondria have retained a small extra-nuclear genome encoding essential components of the electron transport chain (ETC), as well as transfer and ribosomal RNAs required for their translation (Fig. 1a). The ETC is responsible for producing cellular energy through oxidative phosphorylation and maintaining a reducing chemical environment. As such, the genetic information encoded in the mtDNA is essential for maintaining cellular homeostasis. However, due to the absence of several DNA repair pathways, mtDNA exhibits mutation frequencies >100-fold higher than the nuclear genome [1].

**Fig 1.**
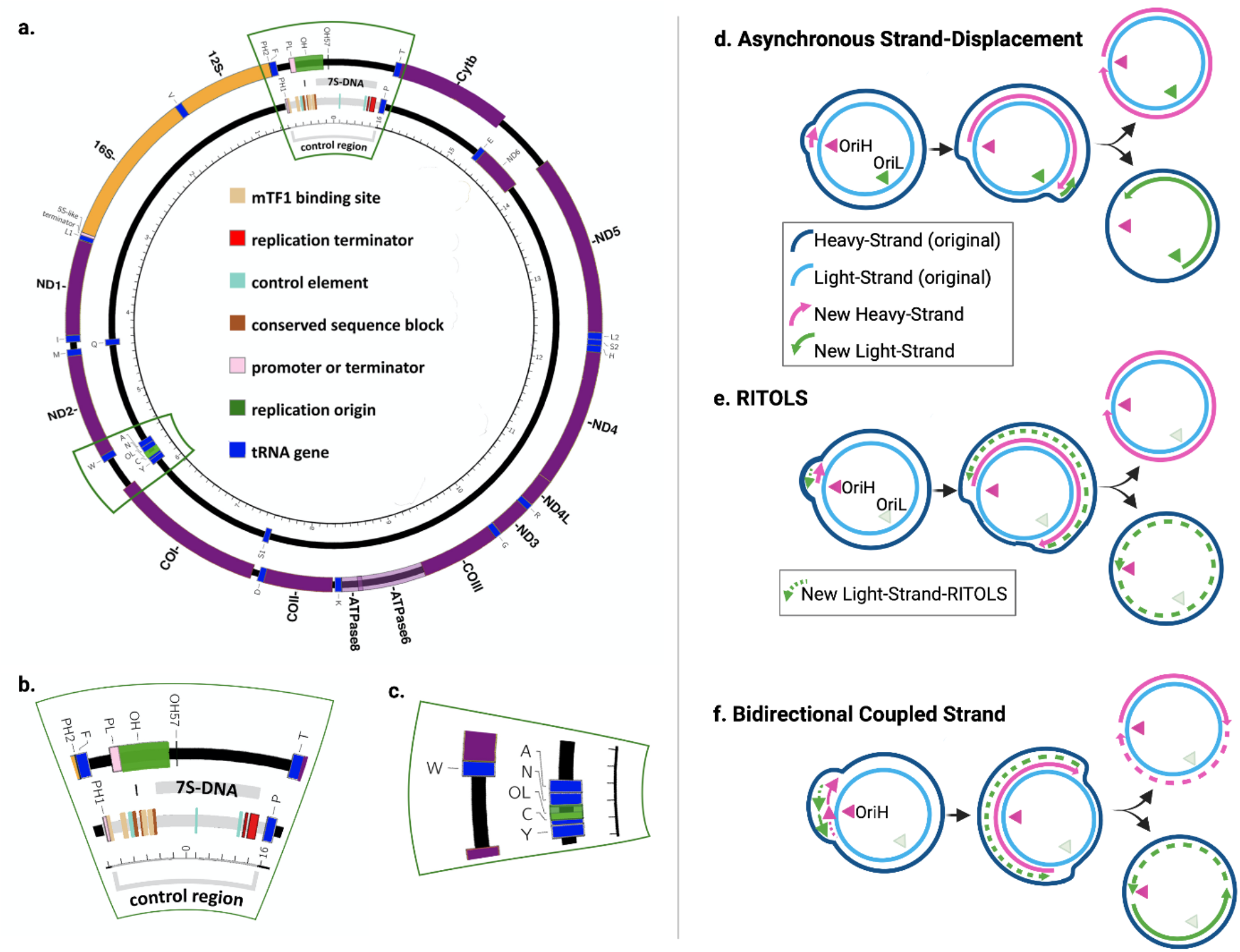
**a** Schematic of mammalian mtDNA and proposed replication models. Gene order and regulatory structures are preserved between humans and mice. Outer ring represents the light strand and the inner ring represents the heavy strand. Genes colored by complex is as follows: *blue*=ribosomal genes; *yellow*=Complex I; *orange*=Complex III; *red*=Complex IV; *purple*=Complex V. **b** Magnified area of the control region. **c** Magnified area of the Ori_L_. **d** Schematic of asynchronous strand-diplacement model as originally proposed by Clayton and colleagues. **e** Schematic of RITOL model. **f** Schematic of strand-synchronous bidirectional replication model. Figure adapted from Lujan *et al*. and licensed under the CC BY 4.0 [61].

Mutations in the mtDNA cause a number of devastating maternally inherited diseases, while the accumulation of mutations in the soma is linked to common diseases of the elderly, including cancer, diabetes, and neurodegenerative diseases (Reviewed in [2]). While an important driver of human health, the mutagenic processes that give rise to these mutations are under considerable debate [3,4]. As originally posited by Denham Harmon, the proximity of mtDNA to the ETC should result in high levels of oxidative damage (*i*.*e*. 8-oxo-dG), yielding predominantly G→T/C→A transversions [5,6]. Counter to this prediction, low levels of G→T/C→A mutations and a preponderance of G→A/C→T and T→C/A→G transitions are observed [7–10]. The presence of these mutations has been interpreted as arising from either base selection errors by DNA polymerase γ (Pol-γ) or spontaneous deamination of deoxycytidine and deoxyadenosine, and not due to reactive oxygen species (ROS) induced 8-oxo-dG adducts.

Regardless of the specific source of mutagenesis, the replication of the mtDNA by Pol-γ is required for fixation of mutations into the genome. Thus, the distribution of mutations can provide clues to the mechanism by which genome replication gives rise to *de novo* mutations. The mechanism of mtDNA replication remains poorly understood, but, in vertebrates, is generally thought to occur via an asynchronous strand displacement mechanism involving two separate, strand-specific, origins [11,12](Fig. 1a-c). In this model, replication is initiated at the heavy-strand (H-strand) origin (Ori_H_), located in the non-coding CR, using a displacement loop (D-loop) as the replicon primer (Fig. 1b,d). Synthesis of the nascent H-strand displaces the original H-strand into a single-stranded state. Upon traversing the light-strand (L-strand) origin (Ori_L_), located approximately 11,000 bp away from the CR, a second replication fork is established and proceeds in the opposite direction, resulting in the original H-strand becoming double-stranded (Fig. 1c,d). Replication is completed when both replication forks complete their circumnavigation. Alternative vertebrate models have been proposed whereby the displaced H-strand is annealed to RNA transcripts, termed RITOLS, that serve to prevent the single-stranded state and act as intermittent priming sites for L-strand replication (Fig. 1e) [13,14]. Visualization of replication intermediates by 2D-gel electrophoresis has also indicated the presence of coupled-strand synthesis involving a more conventional leading/lagging strand replication fork initiating from a bidirectional origin (Ori_b_) in the mtDNA CR or potentially throughout a multi-kilobase “initiation zone” (Fig. 1f) [15–17]. The asynchronous and coupled-strand mechanisms have been proposed to be present at the same time, contingent on the physiological state of the cell [15,18]. Lastly, alternative tRNA genes outside of the Ori_L_ tRNA cluster have been proposed to act as alternative L-strand origins [19,20].

Each of these replication models have significant implications for mtDNA mutagenesis. As hypothesized in previous phylogenetic studies, an asymmetric mechanism of mtDNA replication could explain the phenomena of G/C strand bias, A/T-skew, and mutational gradients seen across taxa [21– 23]. Specifically, the long-lasting “naked” ssDNA replication intermediate in the original model predicts elevated levels of G→A/C→T mutations when the template dC is in the (single-stranded) H-strand due to cytidine exhibiting substantially increased deamination rates when present in a single-stranded state [24]. In this case, the mutational pressure is away from dC content in the H-strand and towards increased dT content. Moreover, genes closer to the Ori_H_ are expected to be more mutation prone than those farther away due to longer times in the single-stranded state. In contrast, both conventional leading/lagging-strand synthesis and intermittent priming models could produce G/C strand bias and/or A/T-skew arising from different mutation frequencies between the leading and lagging strands, a phenomenon observed in bacteria and nuclear DNA (nDNA) replication [25,26], but a mutational gradient stemming from deamination events should be weak or absent due to negligible amounts of ssDNA. Using modern sequencing technologies, the strand asymmetry in transitions has been described in somatic mtDNA mutations [7–10]. More recently, high accuracy sequencing of murine oocytes shows a similar bias towards the strand-asymmetric accumulation of transitions, establishing a mechanistic link between the dominant mutagenic process in somatic tissues and what is seen in population genetics [27]. However, to date, no mutational gradient has been reported outside the context of phylogenetic analyses and it remains an open question if it is an active process or a byproduct of selective pressure over time.

In this report, we have taken advantage of several large high accuracy mtDNA mutation data sets previously generated with Duplex Sequencing (Duplex-Seq) to examine the distribution of somatic mutations in the mtDNA of mice and humans (Sanchez-Contreras & Sweetwyne et al., *in preparation* and [10,28,29]). We find that G→A and T→C transitions, but not their complementary mutations, exhibit a strand-asymmetric gradient delimited by the Ori_L_ and the CR. This gradient is evolutionarily conserved between mouse and humans. The CR also exhibits a remarkably different mutational pattern compared to the coding portion of the genome and is consistent with the presence of a stable D-loop structure bounded by highly conserved regulatory sequence blocks (Fig. 1b). Comparison of the somatic mutational gradient to the distribution of SNPs in the human population, as well as the distribution of bases along the genome across species, shows remarkable concordance. Taken together, our findings demonstrate that an active mutational gradient drives the unequal accumulation of mutations in mtDNA and is most consistent with a strand-asymmetric replication model with an extensive ssDNA replication intermediate. Moreover, this unusual mutagenic process influences population level haplotypes and likely drives genome composition over evolutionary time scales.

## Results

As part of a comprehensive analysis on the effects of aging and mitochondrial-targeted interventions on somatic mtDNA mutation accumulation in eight different mouse tissues, we used Duplex-Seq to collect 34,113 independent, high accuracy, somatic mutations spread across the entirety of the mtDNA molecule (Additional File 1: Supplemental Data 1). In the course of initially analyzing our data, we noted significant variability in the per gene mutation frequency (when looking at individual mutation types). Ordering the genes by their location in the genome, instead of grouping by complex, showed an increasing frequency in G→A mutations, reminiscent of what has been observed in phylogenetic studies (Additional File 2: Supplemental Fig. 1) [23,30]. Intrigued by this observation, we took advantage of the large number of mutations to obtain a higher resolution understanding of how mutagenesis varies across the mtDNA. However, variants are spread out across 85 individual samples with a mean of 401 (range: 48-1496) mutations per sample, corresponding to a mean density of 0.025 mutations/bp. Because mutations are spread across 12 different mutation classes, the mutation density of individual samples would not provide a higher resolution than at a per gene level. To overcome this issue, we combined the data from all tissues to produce the most robust data set possible. Specifically, we divided the genome into 100bp bins (total of 163) and, for each mutation class (*i*.*e*. G→A, G→C, G→T, *etc*), summed the mutation counts observed in each bin across our all samples, separated by age cohort (young (n=40): 4.5 months; old (n=45): 26 months). We then normalized for both genome base composition and variability in sequencing depth of each bin by dividing the mutation count by the total number of wild-type mutable bases sequenced across the constituent samples (Additional File 1: Supplementary Data 2 & 3). This effectively gives a weighted mean of the mutation frequency for each bin for all samples.

By plotting the mutation frequency by genome position (*i*.*e*. bin) in our 26 month old cohort (25,020 mutations), an apparent discontinuous gradient bounded by the Ori_L_ and CR is observed for G→A and T→C transitions, but not their respective complementary mutation types (Fig. 2a,b). An exception is the CR (bins 154-163; genome positions 15400-16,299) which exhibits a notable spike in C→A, but a decline in G→A mutations, whereas both T→C and A→G mutations show increases in the CR, consistent with previous reports [7,27]. Performing separate regressions of the minor and major arcs (bins 1-48 and 53-154 or genome positions 1-4800 and 5,300-15,400, respectively) show highly significant increases in mutation frequency across their respective genomic coordinates (minor arc: G→A slope=8.24±3.06×10^−8^, p=0.007; T→C slope=1.08±0.35×10^−8^, p=0.002; major arc: G→A slope=7.42±0.75×10^−8^, p=5.35×10^−23^, T→C slope=1.23±0.10±10^−8^, p=1.33×10^−35^). With the exception of G→C mutations in the major arc, no other mutation types exhibited a gradient (Additional File 2:

**Fig 2.**
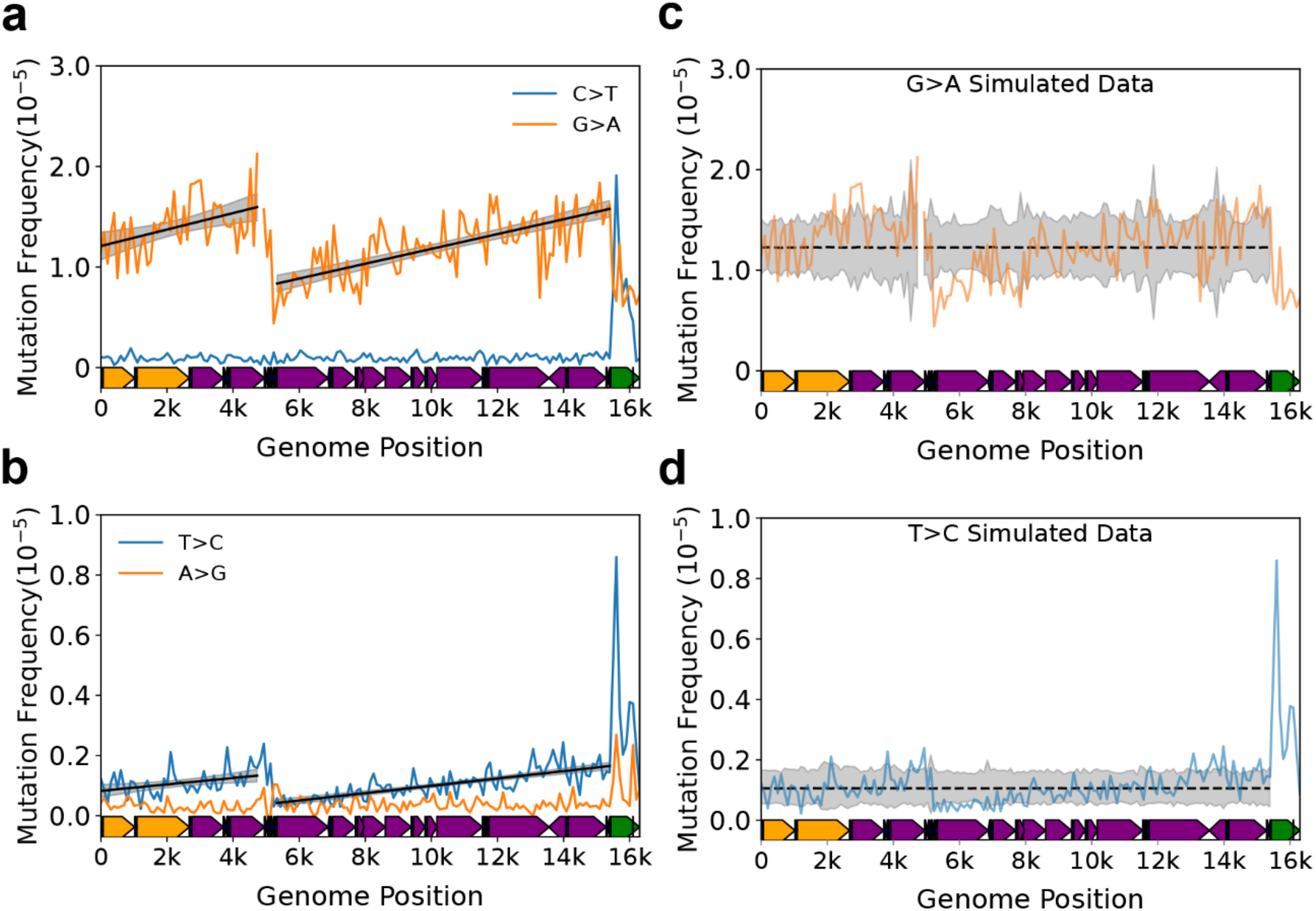
Somatic transitions mutations exhibit a mutational gradient in mouse mtDNA. **a-b** Plot and linear regression (*black line*) of reference strand (*i*.*e*. L-strand) C→T/G→A and T→C/A→G mutation frequencies as a function of genome position. Each data point denotes a 100bp bin. **c-d** Distribution of simulated mutation frequencies of G→A and T→C mutations along the mouse mtDNA. Simulations are based on the data in **a** and **b**. *dotted black line*=bin specific mean; *grey shading=*empirical 95% confidence interval; *red line*=fitted regression, dotted denotes p>0.05 for slope and solid denotes p≤0.05; *red shading*=95% confidence interval of linear regression. Mouse mtDNA structure and coordinates are shown on the x-axis and are the same for all panels (*orange*=rRNA gene, *purple*=protein coding, *dark blue*=tRNA gene, *green*=control region).

Supplementary Fig. 2 and Additional File 3: Supplementary Table 1). Notably, the G→C gradient is >10-fold smaller than the transition-based gradients and its relevance to mitochondrial biology, if any, is unclear. The strand bias (reference L-strand G→A and T→C mutations are equivalent to anti-reference H-strand C→T and A→G mutations, respectively) is consistent with the previous reports in somatic mtDNA mutations and the gradient is most consistent with the previously hypothesized strand-asynchronous replication mechanism with a deamination prone single-stranded replication intermediate involving only two origins of replication [7,11,12,27].

Our mouse mtDNA data set combined mutation profiles of 8 unique tissue types from 6 organ systems, therefore we sought to validate our analysis by accounting for the tissue-specific effects and local differences in sequence contexts identified in these data (Sanchez-Contreras & Sweetwyne et al., *in preparation*). To address the possibility that one tissue type in our data was driving the observed gradient, we performed a leave-one-out approach by eliminating one tissue and then performed the same analysis on the reduced data set, repeating this analysis for each tissue type. As expected, the removal of data of any one tissue type did not alter our findings (Additional File 3: Supplemental Table 2). These results point to the gradient not being an artifact of any single tissue type in our data. We next addressed the potential impact of different local sequence contexts within each bin by performing Monte-Carlo simulations that randomly redistributed each mutation observed in the 153 bins corresponding to the non-CR portion of the genome (genome positions 1-15,400) using a weighted probability for each bin based on its base composition. After redistribution, the mutation frequency was then recalculated for each bin and the procedure repeated 10,000 times. As expected, we observe no gradient in either the major or minor arcs (Fig. 2c,d; Additional File 2: Supplemental Fig. 3; Additional File 3: Supplemental Table 3). Our analysis confirms that the strong positive mutational gradients in G→A and T→C transitions in both the major and minor arcs of the mouse mtDNA are not artifacts and is most consistent with a strand asynchronous replication mechanism and inconsistent with a conventional leading/lagging strand mechanism.

**Fig 3.**
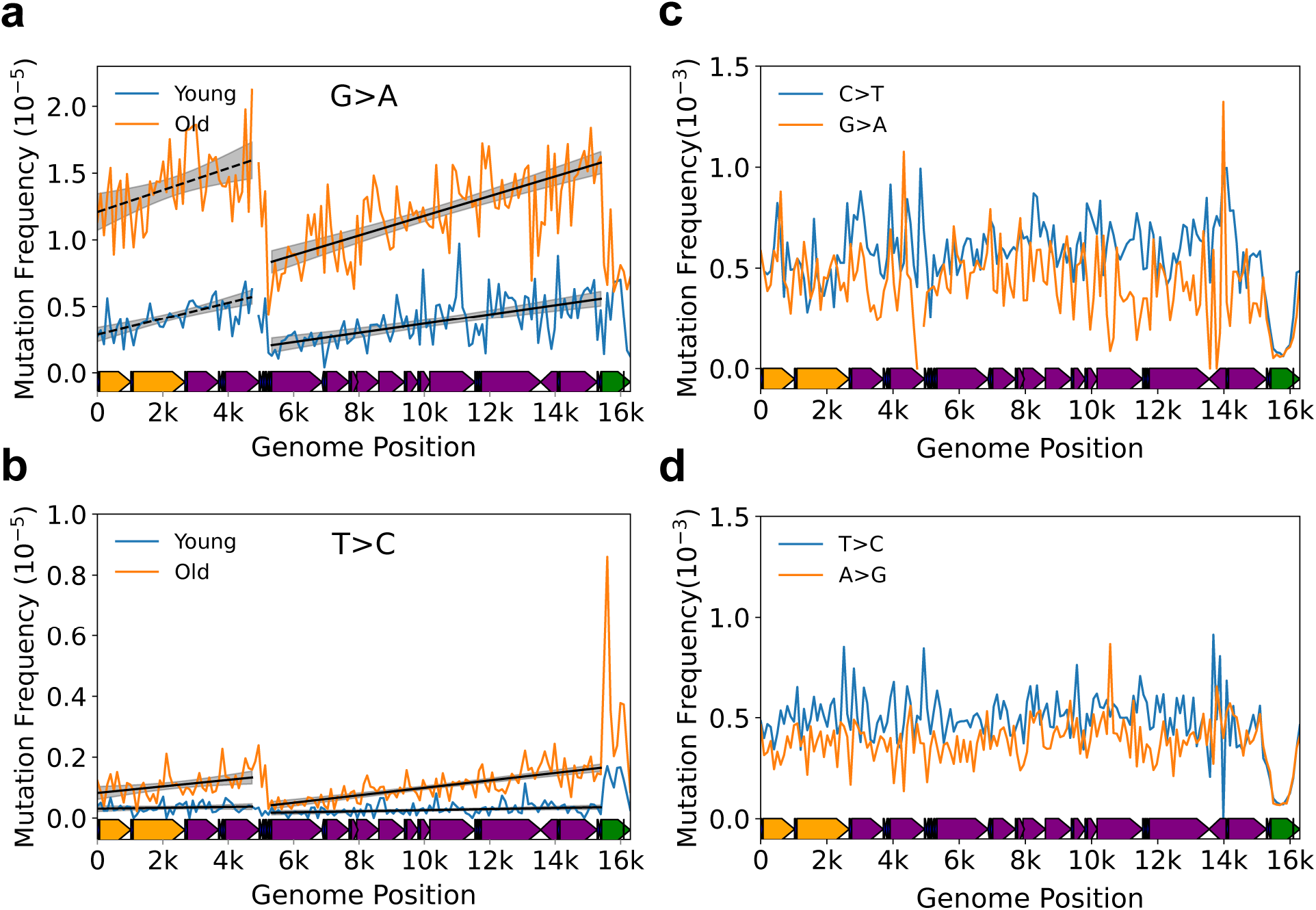
A mutational gradient is established over the course of natural aging and is not directly caused by polymerase-γ base selectivity. **a-b** Changes in the gradient slope of the major arc between 4-6 month old (*blue*) and 26-month old (*orange*) mice. *Black line* and *grey shading*=fitted regression and 95% confidence interval. **c-d** Plot and linear regression (*black line*) of reference strand (*i*.*e*. L-strand) C→T/G→A and T→C/A→G mutation frequencies show an absence of mutational gradient in Pol-γ^exo-^ mice, which lack a functional exonuclease activity in DNA Polymerase γ. Mouse mtDNA structure and coordinates are shown on the x-axis and are as follows: *orange*=rRNA gene, *purple*=protein coding, *dark blue*=tRNA gene, *green*=control region.

### Effects of age and Pol-γ fidelity on the mutational gradient support an asynchronous replication model

A key question is the identity of the biological process giving rise to the observed gradient. As noted previously, the classic asynchronous replication model hypothesizes a long-lived ssDNA intermediate (Fig. 1c). The consequence of this model is that the portions of the mtDNA closest to their initiating origin should disproportionately accumulate G→A and T→C L-strand mutations during the aging process due to more time in the single-stranded state and should manifest as an increase in the gradient slope over time. To test this hypothesis, we made use of the young (4-5 mo; n=9,093 mutations) and old age (26 mo; n=25,020 mutations) cohorts in our data set to evaluate the interaction between aging and genome position on the gradient slope. Both major arc G→A and T→C L-strand gradients, as well as T→C mutations in the minor arc, exhibit a significant increase in their respective slopes during aging (Major arc: G→A interaction=4.21±0.80×10^−8^, p=1.54×10^−7^; T→C interaction=1.03±0.11×10^−8^; p=1.31×10^−21^; Minor arc: T→C interaction=8.38±3.89×10^−9^, p=0.031) (Fig. 3a,b; Additional File 3: Supplemental Table 4; Additional File 1: Supplemental Data 2 & 3). These findings, again, point to the asynchronous replication model as being most consistent with a deamination prone replication intermediate that experiences increased time in the single-stranded state. Furthermore, they demonstrate that this mutational gradient process is the primary driver of age-associated somatic mutations in mtDNA.

While the non-uniform increase in mutations with age is most consistent with deamination, it is possible that some other aspect of mtDNA replication could lead to this pattern. For example, Pol-γ is thought to exist both with and without its p55 accessory subunit, which has been reported to affect fidelity [31,32]. To test the effects of Pol-γ fidelity on the mutation gradient, we reanalyzed the distribution of 30,264 independent mutations obtained from a previous study using Duplex-Seq on mtDNA from mice homozygous for exonuclease deficient Pol-γ (Pol-γ^exo-^) [29]. The loss of exonuclease activity in these mice results in a ∼100-fold increase in mtDNA mutations [29,33]. If the mutational gradient is a fundamental aspect of Pol-γ base selectivity (regardless of the specific cause), we would expect the gradient to still be present or exacerbated in the absence of exonuclease activity. In contrast, if the gradient is due to a non-polymerase source, such as DNA damage, then the frequent misincorporation events of the Pol-γ^exo-^ enzyme should result in a more uniform distribution of mutations across the mtDNA with little to no gradient present.

In contrast to our results in wild-type mice, the strong positive gradient in G→A and T→C transitions is no longer present (Fig. 3c,d; Additional File 3: Supplemental Table 5; Additional File 1: Supplemental Data 4). Instead, we note slight, but statistically significant, negative slopes in G→A and T→C transitions, as well as T→A, C→G, and G→T transversions in the major arc and a slight positive slope in minor arc A→T, but the relative effect size is substantially smaller than what is seen in G→A and T→C mutations in wild-type mice and its relevance in mtDNA biology, if any, is unclear (Additional File 2: Supplemental Fig. 4; Additional File 3: Supplemental Table 1&5). We did not evaluate the distribution of mutations in the CR due to the likely presence of concatemers in the Pol-γ^exo-^ mouse CR [34], the effects of which can be seen by the significantly lower mutation frequencies in bins containing this region (Fig. 3c,d; Additional File 2: Supplemental Fig. 4). Taken together, our analysis showing that the gradient unequally changes across the genome with age and the loss of the strong positive gradient from replication by error prone Pol-γ^exo-^ points to a mechanism that is extrinsic to the polymerase itself and is, again, most consistent with a DNA replication intermediate with a long-lived single-stranded state.

**Fig 4.**
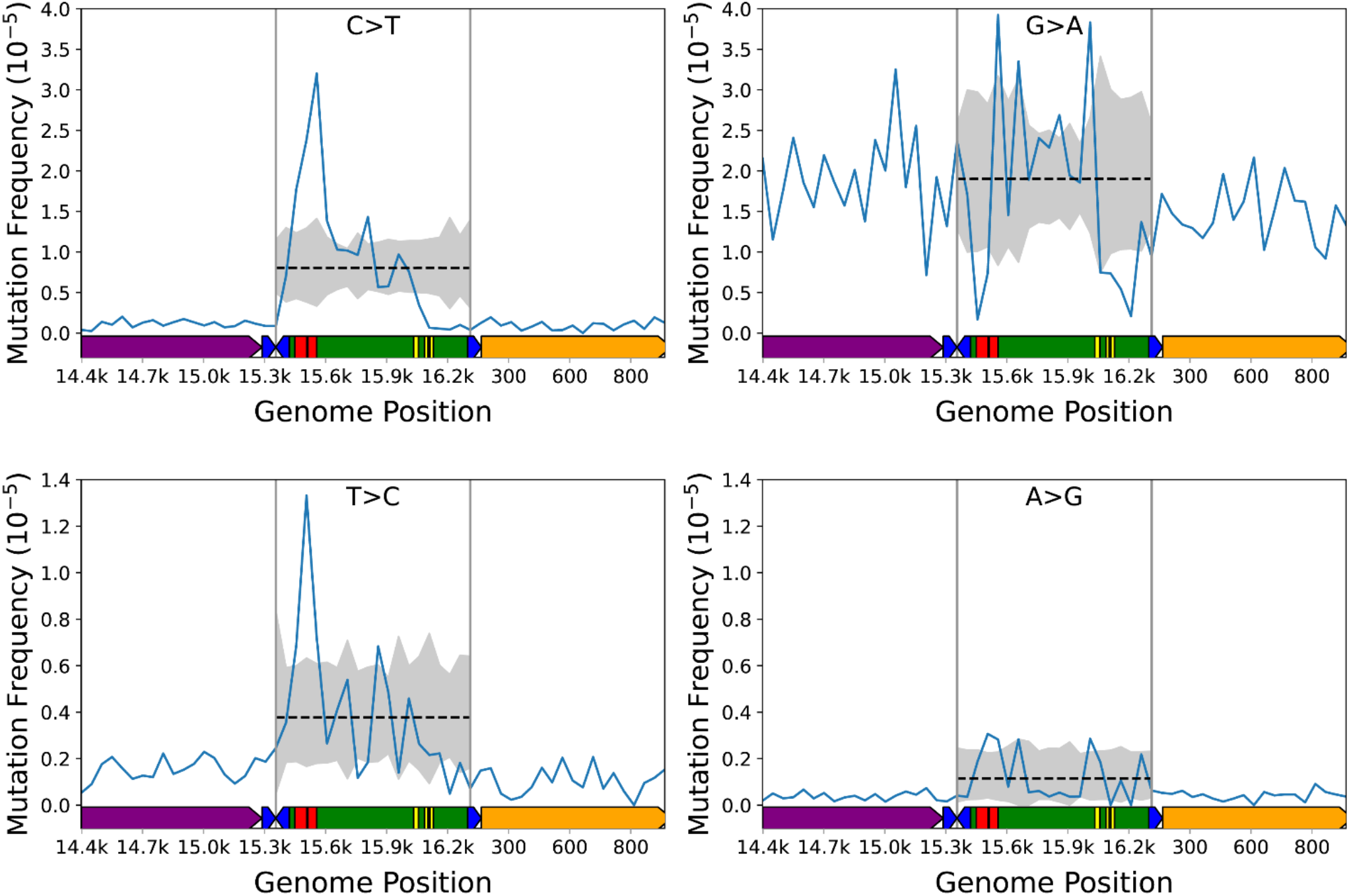
Mutations in the mtDNA control region display a non-uniform distribution and constraints at some loci. The observed per bin mutation frequency (*blue line*) and simulated distribution of data (*dotted black line*=mean, gray shading=experimental 99.975% confidence interval) C→T, G→A, T→C, and A→G mutations. Each data point denotes a 50bp bin. Data points outside the shaded areas are either over- or under-represented compared to random chance. Mouse mtDNA structure and coordinates are shown on the x-axis and are the same for all panels (orange=*mt-Rnr1* gene, *purple*=*Cytb* gene, *dark blue*=tRNA genes, *green*=control region, *red*=ETAS1&2; *yellow*=CSB1-3).

### Conserved regulatory elements exhibit mutagenic ‘hot-spots’ and ‘cold-spots’

The CR contains several important regulatory elements, including both transcriptional promoters, the Ori_H_, several highly conserved sequence blocks (CSB), and extended termination-associated sequences (ETAS), whose specific regulatory functions are incompletely understood (Reviewed in [35]) (Fig. 1b). We and others have noted a distinctly different mutation frequency and spectrum in the CR compared to the coding portion of the genome in both humans and mice [7,27,36], suggesting that the unique function and structure may strongly influence CR mutagenesis, but high-resolution mapping of mutations has not been reported.

The CR lies at the extreme 3’ terminal end of the *M. musculus* mtDNA reference genome, which presents issues during data alignment that gives rise to significant biases in sequence depth and mutation calls. To address this potential bias, we modified the mtDNA reference to place the CR in the middle of the sequence and realigned our data to this modified reference. In addition, we decreased our bin size to 50bp to allow for a higher resolution mapping of mutations. The CR exhibits prominent spikes and troughs that closely correspond to the ETAS, 7s DNA D-loop, CSBs, and the transcriptional promoters (Fig. 4; Additional File 1: Supplemental Data 5) [37]. To confirm our findings, and to determine if any of these conserved sequences comprise a mutational ‘hot-spot’ or ‘cold-spot’, we performed Monte-Carlo simulations using the same strategy as described for our mutational gradient analysis, but with 50bp bins, repeating the sampling 100,000 times, and setting the two-tailed Bonferroni corrected significance to p<0.0025 (Fig. 4; Additional File 2: Supplemental Fig. 5, *black line & grey shading*). The simulations confirm that C→T, T→C, G→A, and A→G, but not other mutation classes, show significant deviations from random sampling in these conserved structures. Of particular interest is a consistent mutational ‘hotspot’ for C→T, T→C, and A→G mutations, but a ‘cold-spot’ for G→A in the ETAS (Fig. 4, *red blocks*). This observation suggests the presence of a structure that is highly prone to certain mutation types and resistant to others or, alternatively, the loss of L-strand dG’s prevents the maintenance of mtDNA. Consistent with the possibility that mutations can be selected against, all four transition types show a significant depletion of variants in the region between CSB3 and mt-tRNA^Phe^ that corresponds to the transcription promoters and mitochondrial transcription factor A (TFAM) binding sites, which are thought to be the source of the Ori_H_ replication primer (Fig. 4, genome position 16,100-16,299)[35]. Interestingly, no high level heteroplasmic or homoplasmic variants have been detected in the same region in human population studies, suggesting that this region is extremely important for mtDNA maintenance [36]. Lastly, all four transitions exhibit a significant spike in the region between CSB1 and genome position ∼15,900 consistent with this region harboring the 7s DNA/RNA D-loop. Taken together, our high resolution analysis of CR mutations highlights the presence of both mutagenic hot-spots and cold-spots that correspond to highly conserved regulatory elements responsible for the distinctive mutational bias previously noted in the CR. Additionally, our data suggest the presence of unique DNA structures within these sequence blocks that differently affect DNA damage and/or replication fidelity and also suggest that some regions important for mtDNA replication may poorly tolerate mutagenesis.

**Fig 5.**
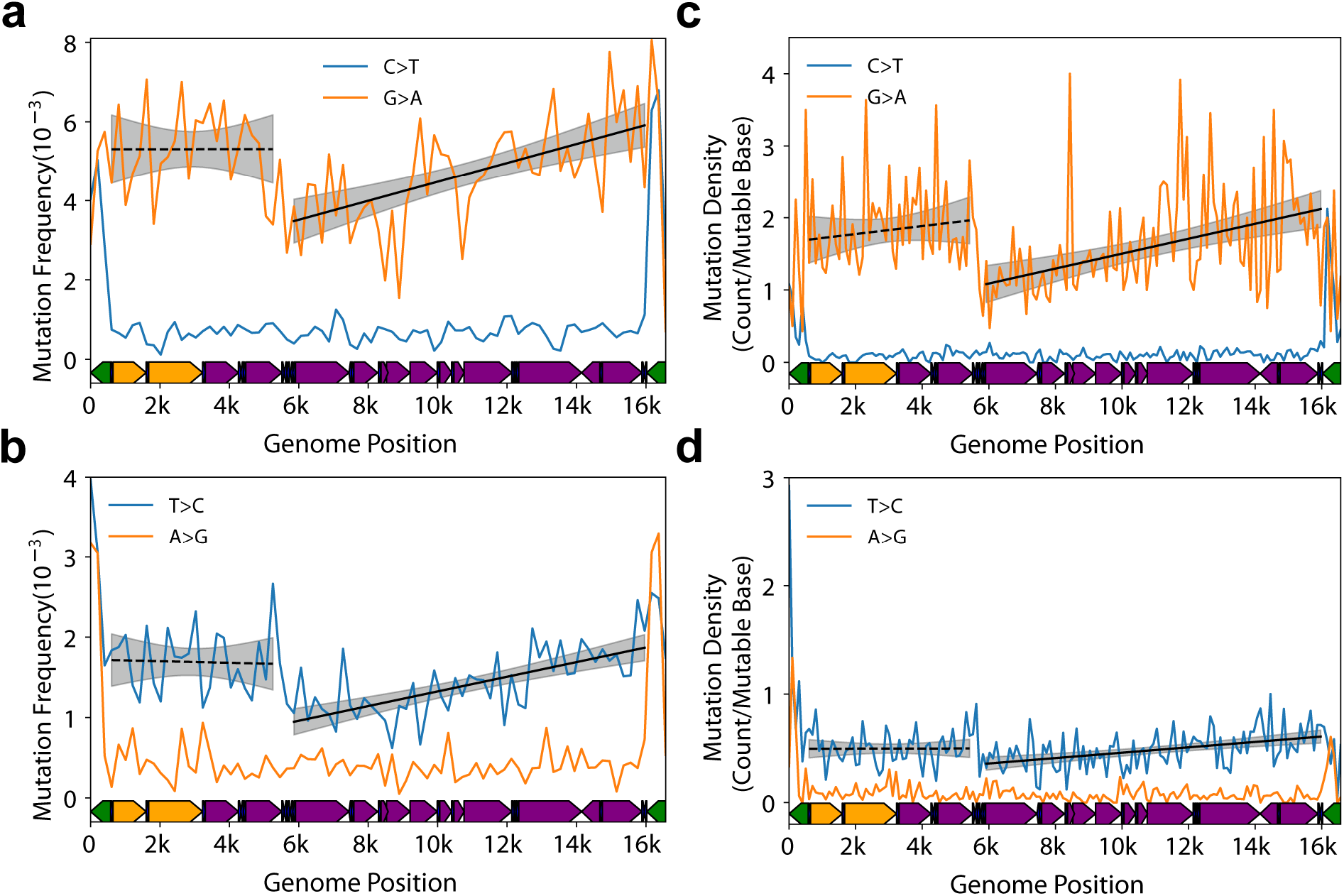
Somatic mutational gradient is conserved in human mtDNA. **a-b** Plot and linear regression (*black line*) of reference strand (*i*.*e*. L-strand) C→T/G→A and T→C/A→G mutation frequencies as a function of genome position from prior published Duplex-Seq data. Each data point denotes a 200bp bin. **c-d** Plot and linear regression (*black line*) of reference strand (*i*.*e*. L-strand) C→T/G→A and T→C/A→G mutation densities as a function of genome position in human tumor data from PCAWG dataset. Each data point denotes a 100bp bin. Human mtDNA structure and coordinates are shown on the x-axis and are the same for all panels and are as follows: *orange*=rRNA gene, *purple*=protein coding, *dark blue*=tRNA gene, *green*=control region. *Black dotted line*= linear regression slope p>0.05; *dashed black line*=linear regression slope p<0.05; *grey shading*= regression 95% confidence interval.

### A mutational gradient is conserved in human mtDNA

We next determined the evolutionary conservation of the patterns we observe in our mouse data. To do so, we made use of prior reported Duplex-Seq data sets for human mtDNA [10,28]. As with the mouse data, we performed a binned mutation frequency analysis with bin size of 200bp due to the reduced number of mutations compared to our mouse data. Consistent with our mouse data, we observe a gradient for both G→A and T→C mutations in the major arc (G→A: slope=4.86±0.93×10^−7^, p=1.93×10^−7^; T→C: slope=1.84±0.26×10^−7^, p=1.69×10^−12^) that is bounded by the Ori_L_ and CR (Figure 4a,b; Additional File 3: Supplemental Table 6; Additional File 1: Supplemental Data 6). Unlike the mouse data, the minor arc did not exhibit an apparent gradient and no other mutation types exhibited a significant increase in either the major or minor arcs (Additional File 2: Supplemental Fig. 6; Additional File 3: Supplemental Table 6).

**Fig 6.**
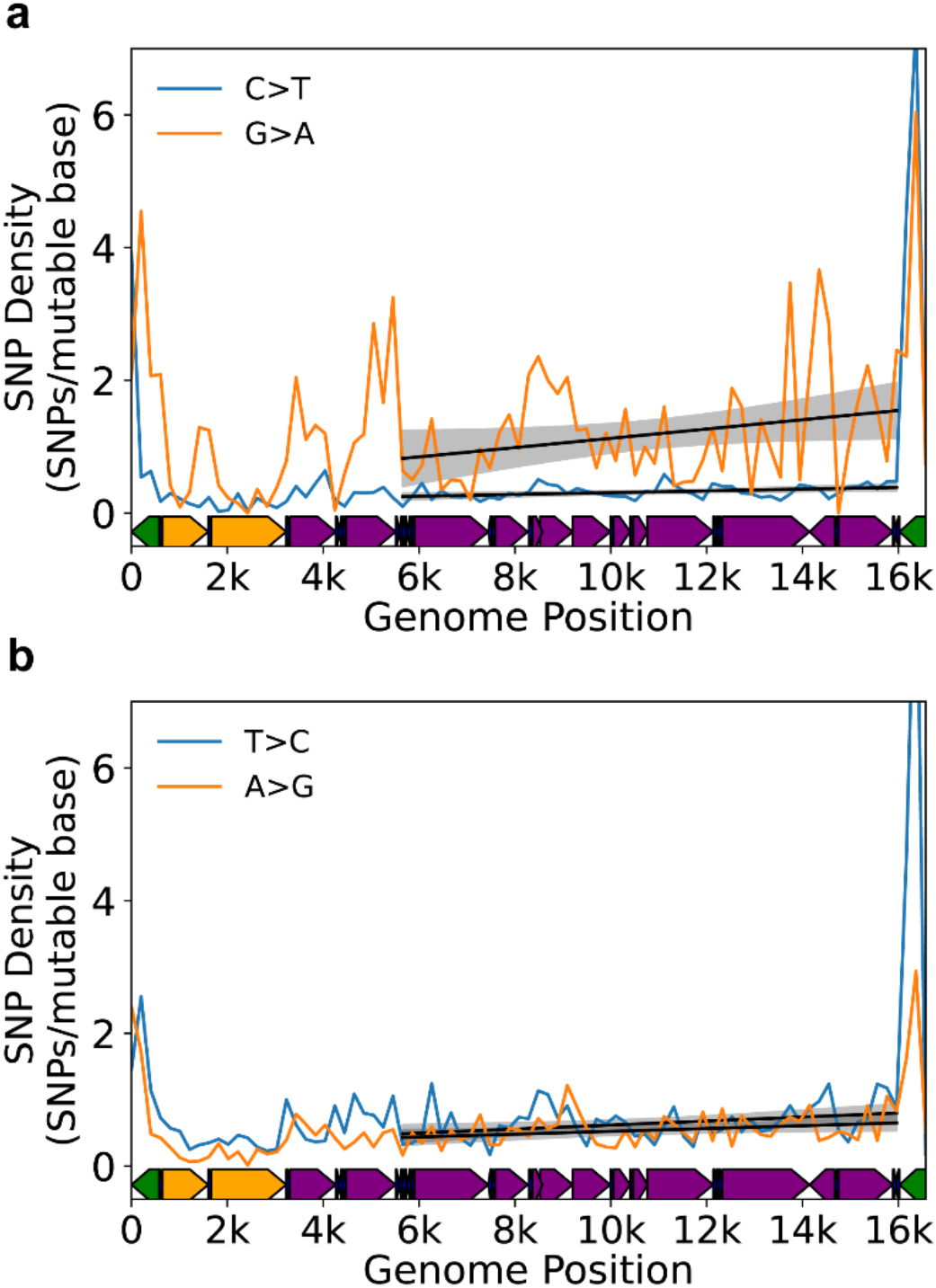
Mutational gradient is detected in major arc in human population SNPs. **a** Density of C→T and G→A SNPs on the L-strand. **b** Density of T→C and A→G SNPs on the L-strand. Data are from Gu *et al*. [40]. *Black dotted line*= linear regression slope p>0.05; *dashed black line*=linear regression slope p<0.05; *grey shading*= regression 95% confidence interval. Human mtDNA structure and coordinates are shown on the x-axis and are the same for all panels and are as follows: *orange*=rRNA gene, *purple*=protein coding, *dark blue*=tRNA gene, *green*=control region.

Our analysis points to a somatic mutational gradient as an evolutionarily conserved feature of vertebrate mtDNA. However, all our data were collected using Duplex-Seq, leaving open the possibility that the gradient pattern is an artifact of our Duplex-Seq protocol or our data analysis pipeline. While we consider this scenario unlikely, we sought to observe this gradient in an independently generated data using more conventional sequencing approaches. Somatic mtDNA mutations occur at very low frequencies (∼10^−6^-10^−5^), making their detection with conventional sequencing difficult [38]. To overcome this limitation, we analyzed mtDNA mutation call data published by the Pan-Cancer Analysis of Whole Genomes (PCAWG) Consortium [39]. This data set consists of 7,611 independent somatic variants (variant allele fraction (VAF)>0.01; mean VAF=0.2) from 2,536 tumors across 38 different cancer types. Because cancer is a clonal process arising from a single cell, the detected variants are largely a snapshot of the mtDNA mutations present early in tumor formation and have much higher VAFs than what is typically detected in Duplex-Seq data. Importantly for our purpose, this characteristic of the tumor data is expected to largely eliminate the potential confounder of low frequency artifacts giving rise the observed gradient.

We divided the genome into 100bp bins and, for each mutation type, calculated the mutation density (i.e. mean number of detected mutations per wild-type base) in each bin (Additional File 1: Supplemental Data 7). Consistent with our Duplex-Seq data, we observe a clear gradient in both G→A and T→C transitions, but not their complement, that increases along the major arc (G→A p=7.43×10^−8^; T→C p=1.16×10^−5^)(Fig. 5c,d; Additional File 3: Supplemental Table 7). Both T→A and C→G transversions report a negative slope in the major arc and C→T and G→T exhibit a positive slope in the minor arc, but the magnitudes are extremely small and are likely a regression artifact. No other mutation types exhibit a gradient (Additional File 2: Supplemental Fig. 7; Additional File 3: Supplemental Table 7). These data confirm both the presence of a mutational gradient in the major arc and that our results are unlikely to be due to an unknown issue with Duplex-Seq. Taken together, both our Duplex-Seq data and the PCAWG data recapitulate our findings in mouse mtDNA, pointing to the strong evolutionary conservation of G→A and T→C gradients among vertebrate species.

**Fig 7.**
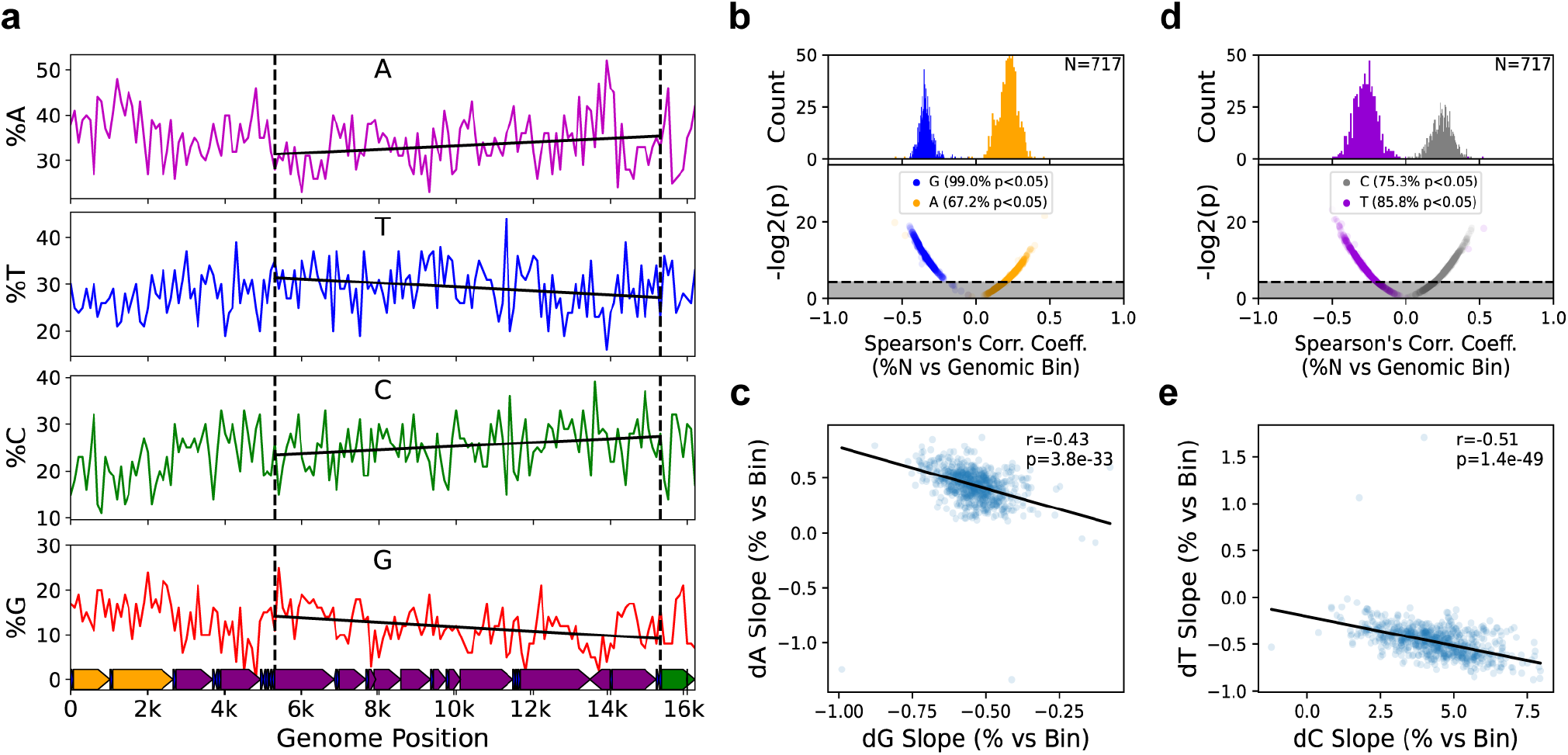
Genome composition bias mirrors the somatic gradient in mammals. **a** Base composition gradient as exemplified by the murine mtDNA. Vertical dashed lines delimit the major arc. Solid black lines are the best fit regression by robust linear regression. Slopes are significantly different from zero in all cases. Gene coloring: *orange*=rRNA gene, *purple*=protein coding, *dark blue*=tRNA gene, *green*=control region. **b**,**c** Anticorrelation of dG and dA base composition in mammals. Most mammalian genomes show a statistically significant spatially dependent depletion of dG and enrichment of dA in the major arc and the strength of dG depletion is directly proportional to dA enrichment in a species dependent manner. **c**,**d** dT and dC show a similar anti-correlative pattern as dG and dA.

### A mutational gradient is mirrored in germline SNPs and genome base composition

Previous work has noted similarities in the strand orientation and simple mutational spectra between somatic mtDNA mutations and population level SNPs, suggesting a similar causative driver of population level mtDNA sequence diversity [7,9,27]. We sought to further explore this relationship by determining if the mutation gradient is reflected in the distribution of inherited single nucleotide variants, as would be expected if this process is active in the germline. We initially sought to test this hypothesis by mapping mutations obtained with Duplex-Seq of mouse oocytes [27], but the total number of mutations (N=691) was insufficient to detect a gradient. We next evaluated the distribution of homoplasmic SNPs in the human mtDNA by downloading a recently published list of 44,494 SNPs obtained from MITOMAP and phylogenetically corrected such that each SNP was likely the result of an independent *de novo* event [40,41]. Using the same binning approach as our human somatic data, we calculated the mutation density (*i*.*e*. number of *de novo* SNPs per mutable base) in each bin. For this analysis, we limited our analysis to the major arc due to 1) the absence of a clear minor arc gradient in our human somatic data and 2) evidence of regions with an underrepresentation of SNPs in rDNA genes [41]. Consistent with our somatic data, we observe a significant positive gradient in G→A and T→C (Fig. 6a,b; Additional File 2: Supplemental Fig. 8; Additional File 3: Supplemental Table 8). Notably, complement SNP types (C→T and A→G, respectively), as well as G→C SNP, show significant gradients, but the magnitude of their slope, especially relative to G→A SNPs is substantially smaller. We sought to further validate this observation by performing this same analysis on a recently reported database of homoplasmic SNPs from 196,983 individuals [41]. As with our initial data set, we observe a significant correlation between SNP density and genome position of G→A SNPs (ρ=0.26; p=0.046; Spearman correlation). We also observe a significant correlation between G→C SNP and genome position (ρ=0.307; p=0.019; Spearman correlation) similar to the MITOMAP based dataset. No other significant correlations were observed (Additional File 3: Supplemental Table 9). Thus, we are able to confirm that, at the very least, a G→A gradient is present in human polymorphisms, consistent with our somatic mtDNA data, and further supports the idea that the mechanism of mutagenesis in the somatic tissue is likely the direct driver of human mtDNA variation.

The strong conservation of the somatic gradient between mice and humans and the presence of the gradient in human SNP data suggest that this unusual mutational pressure is likely a major driver of sequence diversity across species. Our somatic data point to a sustained G→A and T→C mutational pressure of the L-strand with relatively little reversion. Over the long term, the L-strand is expected to exhibit a spatially dependent depletion of dG and dT bases along the major arc and a concomitant increase in dA and dC bases until some selective equilibrium is reached (Fig. 7a). Phylogenetic analyses on the sequence differences between related species such as primates has been shown to exhibit a gradient effect in T→C transitions [30]. Analysis of a relatively small number of vertebrate species (N=118) has also suggested that this phenomenon is likely a general aspect of vertebrate mtDNA biology [23].

We sought determine the generality of the gradient phenomenon by expanding these findings to include the significantly increased number of vertebrate mtDNA sequences now available (N=3,614). Performing this analysis on all available mammalian mtDNA sequences in the NCBI RefSeq database (N=717) shows that the majority of sequences exhibit a significant spatially dependent depletion of dG and dC (*i*.*e*. negative correlation coefficient) and a similar enrichment (*i*.*e*. positive correlation coefficient) in dC and dT(Fig. 7b,d), confirming that this is a general phenomenon in mammalian mtDNA. While consistent with our hypothesis, the correlation 1) does not inform on the magnitude of the correlation and 2) does not explicitly link the change in the abundance of one base type with another. Specifically, the magnitudes of the dG and dT composition slopes should be anti-correlated with the respective dA and dC slope magnitudes within the same species. As can be seen in Figure 7c & 7e, with a few exceptions, the slopes of dG and dA content, as well as dT and dC content, are strongly anti-correlated (dG/dA Spearman’s ρ=0.43, p=3.8×10^−33^; dT/dC Spearman’s ρ=0.51, p=1.4×10^−49^) across currently available mammalian mtDNA sequences with the direction of the anti-correlation consistent with a graduated G→A and T→C mutation pressure. We next extended this approach to other vertebrate classes, including birds (N=656), reptiles (N=212), and fish (N=2029). We did not evaluate non-vertebrate mtDNA sequences due to higher levels of structural heterogeneity and gene composition in these phyla. Like mammals, the majority of species within each vertebrate class show significant gradients in mtDNA composition that are strongly anti-correlated in their dG/dA content, as well as dT/dC content, indicating that this graduated mutation pressure is highly conserved across widely divergent species that inhabit significantly different ecological niches and are subjected to very different selective pressures (Additional File 2: Supplemental Fig. 9 & 10). Interestingly, several species strongly deviate in either gradient direction and/or correlation strength, suggesting that these species are subject to different selective pressures on their mtDNA (Additional File 2: Supplemental Fig. 9 & 10). Taken together, our data point to the mutational process driving the accumulation mutations in somatic tissues being the likely mechanistic driver of population level polymorphisms and sequence composition in vertebrates.

## Discussion

The advent of ultra-high accuracy sequencing methodologies have opened up the possibility of studying the mutagenic processes in mtDNA in greater detail. Both we and others have used Duplex-Seq, a method with an error background of <1×10^−7^, to study somatic mtDNA mutations [7,10,27– 29,43,44]. These studies have broadly shown that mutations are heavily weighted towards G→A/C→T and T→C/A→G transitions with very low levels of transversions, including the canonical ROS-associated G→T/C→A mutations. In addition, these studies have shown a strong strand bias, with G→A/C→T mutation being more prevalent when the dG base is in the L-strand. A notable difference in the mutational frequency and spectrum in the CR is also reported. While these studies have provided a broad understanding of mtDNA mutagenesis, the very low frequency of mutations (<1×10^−5^) means that, for any given sample, only a few dozen to a few hundred mutations are typically detected, leaving conclusions about how these mutations are distributed unclear beyond broad regional differences (*i*.*e*. CR vs coding or between genes). In this study, we aggregated several pre-existing Duplex-Seq data sets to better asses the distribution of mutations across the mtDNA molecule at significantly higher resolution than what has been previously reported.

In addition to recapitulating previous findings showing a strong bias towards transition mutations over transversions and a higher mutation load in the CR, our analysis shows a strikingly non-uniform gradational distribution of G→A and T→C transitions, but not their complement, along the coding portion of the mtDNA. The totality of our data is most consistent with an asynchronous strand displacement mechanism with a long lived, deamination prone, single-stranded H-strand. A key aspect of our data that supports this hypothesis is the increased slope of G→A and T→C mutations with advancing age. Any alternative replication model without a ssDNA intermediate would need to account for how deamination-linked mutations could disproportionately increase as a function of genome position over time beyond what is present in the gradient. The RITOL and strand-synchronous models lacks any substantial ssDNA, with >80% of the displaced H-strand estimated to be annealed with RNA in the RITOL model (Fig. 1d,e)[13,14]. Our Pol-γ^exo-^ data suggest that the gradient is not due to a simple interaction between mtDNA base composition and polymerase base selectivity.

Holt and colleagues have reported that the synchronous and asynchronous mechanisms can exist simultaneously, with the balance between these two mechanisms the result of the cell’s physiological state [15]. While our data do not support a classic leading/lagging strand mechanism, they do not entirely refute its existence in all cases. Leading/lagging strand synthesis may be part of a stress response pathway to quickly reestablish copy number levels. In support of this possibility, withdrawing mtDNA depleting ethidium bromide from cells results in a burst of mtDNA synthesis with fully double-stranded replication intermediates, which is interpreted as being due to a leading/lagging strand replication fork [1]. Consistent with this idea, modulating the level of the mitochondrial transcripts via changes in Twinkle helicase levels has been reported to switch between strand asynchronous and lead/lagging strand synthesis [18]. However, our data are from tissues of unstressed wild-type animals without known perturbations to mtDNA gene expression, pointing to the asymmetric model being the predominant mtDNA synthesis mechanism under normal physiological conditions.

A lingering question in the field of mtDNA replication concerns the conservation of the mtDNA replication mechanism across taxa. The mitochondrial genome exhibits a wide range of sizes, structures, and noncoding regulatory regions between phyla and kingdoms, suggesting that different replication mechanisms were retained or acquired since the initial endosymbiosis event that gave rise to mitochondria. For example, while vertebrates make use of a relatively compact CR with an initiating origin and distal counter-directional origin, invertebrates tend to make use of a large highly AT-rich region with only one confirmed origin and one likely late-firing proximal counter-directional origin [45]. Plant mtDNA likely uses an entirely different recombination-dependent and/or rolling circle mechanism without clearly defined replication origins [46]. Our data indicate that mapping of somatic mutations provides an alternative approach to mapping origins of replication and other potential regulatory structures that is free of the complications inherent to interpreting 2D-gels and electron micrographs. Indeed, an analogous strategy has been used to map origins of replication in the human genome by taking advantage of ultra-mutated tumors [47].

We can clearly discern the location of the Ori_L_ in both mouse and human data sets. These data also argue against the proposed use of other tRNAs as L-strand priming sites, as well as a large ‘initiation zone’ for replication, as these models predict either multiple discontinuous gradients or lack a gradient entirely. Instead, our data are consistent with a single L-strand origin in mammalian mtDNA. Moreover, with our high-density mouse data set, we mapped areas of mutation over- and under-abundance in the CR that correspond to sequence blocks essential for mtDNA H-strand replication. Significant deviations are not obvious in the regions flanking the CR other than the Ori_L_, as would be expected if other sequences in these areas were essential for intermittent priming. Notably, avian mtDNA lacks the predicted stem-loop structure of the mammalian Ori_L_ and 2D-gels point to initiation sites across the entirety of the mtDNA, providing a potential model system to further investigate these alternative origin models in vertebrates [17]. In line with this idea, we attempted to analyze Duplex-Seq data for similar patterns in non-vertebrate organisms, *D. melanogaster* [43,48] and *A. thaliana* [44], but the number of mutations were too low and the density too sparse to observe a clear signal, leaving this for future work.

Human population studies have previously identified a bias in the occurrence of G→A and T→C SNPs of the L-strand, as has comparison of human mtDNA sequences with those of evolutionary related species[49,50]. We previously noted that this bias mirrors the strand asymmetry seen in somatic mutagenesis of mtDNA through a process that is continuous throughout life and hypothesized that this pattern was consistent with mtDNA replication via an asymmetric model [7]. Our data extend this observation to also include a gradient in SNP distribution along the genome, as well as genome base composition, further strengthening the link between somatic and germline processes. Our analysis in genome composition point to this gradient being largely, but not universally, conserved in vertebrates. The strength of this gradient, as highlighted by the variation in anticorrelation between complementary bases and the presence of species that deviate significantly from the trend line, can vary significantly. An important aspect of these observations is that they provide a feasible opportunity to mechanistically study the processes that give rise to genetic variation at the population and taxonomic levels. This is especially pertinent in species with very high or very low mutation rates or show unusual biases in genetic variability. Indeed, recent work in *A. thaliana* linked mismatch repair with the low mutation rate seen in this species, and three species of angiosperm genus *Silene* with notably different mutation rates showed corresponding patterns in somatic mtDNA mutations [44,51], lending credence to the idea that studying somatic mutagenic processes can inform on evolutionary and population level patterns. A systematic analysis of somatic mutations in species with unusual genome composition or SNP patterns may provide insight into how and why these species deviate so significantly from related species and clues to their natural history.

## Conclusions

The growing quantities of high-accuracy sequencing data generated from such technologies as Duplex Sequencing has provided the ability to elucidate several mutagenic patterns and biases previously unobserved at the somatic level. Taken together, these patterns argue for an asymmetric strand-displacement replication model, as originally posited by Clayton and colleagues [11,12] and against a more conventional leading/lagging strand replication fork. The mutations that arise as a consequence of genome replication are likely from deamination events of the single-stranded intermediate which affect genetic variation and, ultimately, genome composition.

## Methods and Materials

### Duplex Sequencing Data and Processing

The DNA libraries and sequencing of the Duplex-Seq libraries was performed as indicated in the original publications. Specifically, the human data was obtained from normal tissue data from Baker et al. (SRA accession PRJNA449763) [10] and Hoesktra et al. (SRA accension PRJNA237667) [28]. Five young and six old wild-type mouse data was generated from male C57Bl/6J at 4-5 and 26 months of age, respectively. Aged mice were obtained from the NIA aged rodent colony at an age of 22-23 months and then housed at the University of Washington animal facility until the desired age under approved conditions. Animals were euthanized at the indicated age and a ∼2 mm section of the heart (apex), liver (lobe VI posterior), kidney (outer cortex), skeletal muscle (proximal gastrocnemius), brain (both cerebellum and hippocampus), and eye (retina and eye cup) were flash frozen and stored at -80°C until processed for sequencing. A ∼1mm tissue punch was used to obtain a representative tissue sample from brain regions. Half of the retina was used for processing and eye cups were used in their entirety. Total DNA was purified from each tissue punch/sample using a QIAamp Micro DNA kit using the manufacturer’s protocol. Total DNA was prepared for Duplex Sequencing using our previously published protocol with modifications as described in Hoekstra et al. [28,52]. Pol-γ^exo-^ mouse data was generated from Pickrell et al. (SRA accension PRJNA729056)[29].

After obtaining the raw data, we processed all Duplex-Seq data using v1.1.4 of our in house developed Snakemake-based Duplex-Seq-Pipeline to ensure that all data were uniformly processed with the exception that different data sets had different unique molecular identifier (UMI) and read lengths[53]. Briefly, we perform a reference free consensus approach for error correction similar to that reported in Stoler et al. [54]. The UMI and any associated spacer sequence are parsed from the read, the read 1 and read 2 UMI from a read pair is sorted alphabetically and associated with the read’s SAM record when converted to an unaligned BAM file (See Stoler et al. [54] for details). After the UMIs from read 1 and read 2 are sorted alphabetically such that all reads derived from the same strand of the same parental molecule are grouped together. UMI families from opposite strands of the same parental DNA fragment are grouped sequentially in the sorted file. A per-position consensus is generated for each single-strand is generated with a cut-off of 70% identity and a minimum of 3 reads with the same UMI being required to call a consensus, as previously described [52]. The double-strand consensus is then generated from the two single-strand consensuses sharing the same UMI, if present, with the exception that the identity of the base must match between the two single-strand consensus. Reads with >2% ambiguous bases are removed from further analysis. The resulting post-processed fastq files are aligned against the reference genome (hg38 chrM for the human data and mm10 chrM for mouse the data) using bwa v0.7.17 [56]. The overlapping portions of reads and 10-cycles of the 5’ and 3’ ends of the reads are clipped using fgbio (https://github.com/fulcrumgenomics/fgbio) and adapter sequences are removed using Cutadapt [57]. Insertion/deletion (in/del) realignment and in/del left alignment were performed by the Genome Analysis Toolkit (v3.8.x). A draft list of variants is generated using the pileup functionality of samtools [58]. Reads containing non-SNP variants (defined as a variant allele fraction >40%) are parsed out and subjected to BLAST-based alignment against a database containing potential common contaminants (dog: canFam3; bovine: bosTau9; nematode: ce11; mouse: mm10; human: hg38; rat: Rnor_6.0). The inclusion of our target genome in this database also allows for the identification of pseudogenes. Reads that unambiguously map to the same coordinates as the original alignment are kept and the remaining reads and associated variants are removed from further analysis. An exception to this process was made for mouse mtDNA due to the presence of a ∼5000bp nuclear pseudogene with perfect identity to mm10 chrM [59]. In this case, any ambiguous BLAST alignments mapping to this region were assumed to be mitochondrial in origin and kept. The mutated reads passing our BLAST filter are merged back with the non-mutated reads and mutation frequencies calculated based on the provided target coordinates.

To generate the bin data for each age cohort (*i*.*e*. 4.5mo and 24-26mo), we divided the genome up into the indicated bin size and then calculated the mutation frequency for each bin, *i*, and mutation type, *N* (*i*.*e*. G→A, T→C, *etc*), by 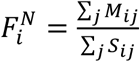, where *M*_*ij*_ is the mutation count of type *N* in bin *i* of sample *j* and *S*_*ij*_ is the number of sequenced bases in bin *i* of sample *j* that are mutable for mutation type *N*. A mutation was only counted if its variant allele fraction (VAF) was <1% to minimize the effects of inherited or early arising mutations.

For the D-loop focused analysis, we generated a new partial mm10 chrM reference consisting of the CR portion with the flanking 1000 bases (chrM 14,400-16299::chrM 1-1001) in order to allow for better alignment of reads across the entirety of the mtDNA. The binning process was performed as described, but with 50bp bins. Per bin mutation frequencies were calculated as described above.

### Tumor sequencing data

Tumor single nucleotide variant data was generated using the methods outlined previously and obtained from The Cancer Mitochondria Atlas data portal (https://ibl.mdanderson.org/tcma/download.html) [39]. Similar to our Duplex-Seq analysis, the human mtDNA was divided into 100bp bins and the number of variants of each of the 12 mutation classes was divided by the number of wild-type base of the respective mutation class within each bin (i.e. #G→A/#G’s, etc).

### SNP and Genome Composition data

The SNP data sets were obtained from Gu *et al* and Bolze *et al* using their respective procedures [41,42]. Briefly, the Gu *et al*. data set is comprised of 44,334 SNPs reported in the MITOMAP database that was filtered for haplotype private variants. The Bolze *et al* data comprises 14,283 homoplasmic variants from 196,324 unrelated individuals. Because this dataset is not limited to haplotype private SNPs, as the Gu *et al*. data, we limited our analysis to rare SNPs (population frequency ≤1:1,000) in order to minimize population structure of the data from confounding our analysis. SNPs occurring more than once were assumed to have arisen from a single *de novo* event. We divided to human mtDNA into 200bp bins and then calculated SNP density by summing the number of variants of each of the 12 mutation classes and then dividing by the number of respective wild-type bases within each bin (i.e. #G→A/#G’s, etc).

For the genome composition analysis, a complete set of curated mtDNA sequences and annotations were downloaded from the NCBI Reference Sequence project (https://ftp.ncbi.nlm.nih.gov/refseq/release/mitochondrion/) in GenBank format. Entries were parsed by taxonomic Class and filtered to separately keep mammals, birds, reptiles, and fish. mtDNA sequence corresponding to the major arc of each species was extracted and divided into 100 bp bins and the nucleotide composition calculated as a percent of each base type. The slope and/or correlation coefficient for the change in genome composition as a function of bin number (*i*.*e*. genome position) for each individual species was then calculated as described below.

### Statistical Analysis

Statistical analysis was performed in python using either Statsmodels (https://github.com/statsmodels/statsmodels) or SciPy [60], where indicated. Linear regression analysis was performed with the Python Statsmodels library using a robust linear model with Huber’s T function as the M-estimator for down-weighting outliers. A robust linear regression model was used due to the violation of the assumptions of normality or homoscedasticity in some data sets that is required for ordinary linear regression models. To establish the effect of aging on the gradient slope, a robust linear regression model with the addition of an interaction term between age and bin number (*Y* = *α* + *β* _*bin*_ * *bin + β*_*bin*_ * *age* + *β*_*binxage*_(*bin* * *age*)),with age being the categorical classifier with value 0 (young) or 1 (old), was used.

Monte-Carlo modeling of random mutagenesis was performed by first dividing the indicated genome interval (*i*.*e*. coding region vs CR) into the indicated bin size. A weighted probability for each mutation type (*i*.*e*. G→A, T→C, *etc*) to occur in each bin was calculated by dividing the cumulative depth of the wild-type mutable base in a bin by the cumulative depth of the wild-type of the same mutable base across the indicated genome interval. For each mutation type, the total number were randomly distributed across the bins using the calculated weights and then a per bin mutation frequency was calculated by dividing the number of mutations distributed in a bin by the cumulative sequencing depth of the mutable base within the same bin. This procedure was repeated 100,000 times for the D-loop analysis and 10,000 times for the coding region gradient analysis. An experimental Bonferonni corrected confidence interval for determining mutational hot-spots and cold-spots was set to 99.75%. Datapoints outside this range were considered significantly different from random chance.

## Author Contributions

Conceptualization: MS-C, MTS, and SRK; Software: BFK and SRK; Sample and Data collection: MS-C, MTS, KAT, MJH, EKS, JF, JAW, MDC; Formal analysis: SRK; Visualizations: MTS and SRK; Writing-initial draft: SRK; Writing-review and editing: MS-C, MTS, BFK, KAT, PSR, DJM, and SRK; Funding acquisition: MTS, PSR and SRK; Supervision: MS-C, MTS, PSR, DJM and SRK; Resources: PSR, DJM, and SRK. All authors read and approved the final manuscript.

## Availability of data and materials

The Duplex-Seq-Pipeline is written in Python and R, but has dependencies written in other languages. The Duplex-Seq-Pipeline software has been tested to run on Linux, Windows WSL1, Windows WSL2, and Apple OSX. The software can be obtained at https://github.com/Kennedy-Lab-UW/Duplex-Seq-Pipeline and https://doi.org/10.5281/zenodo.5084120 under the BSD license. The normal mouse data is available at SRA accension PRJNA727407. Only wild-type, non-intervention samples were used. The Pol-γ^exo-^ mouse data is available at SRA accension PRJNA729056). The human data is available at SRA accession PRJNA449763 and SRA accension PRJNA237667. Only normal control samples were analyzed.

## Ethics approval

Wild-type mouse tissues were collected from mice at the University of Washington under IACUC approved protocols. Previously published human and mouse data sets were collected under the terms described in their respective publications.

