## Supplemental Figures for "A mutational gradient drives somatic mutation accumulation in mitochondrial DNA and influences germline polymorphisms and genome composition"

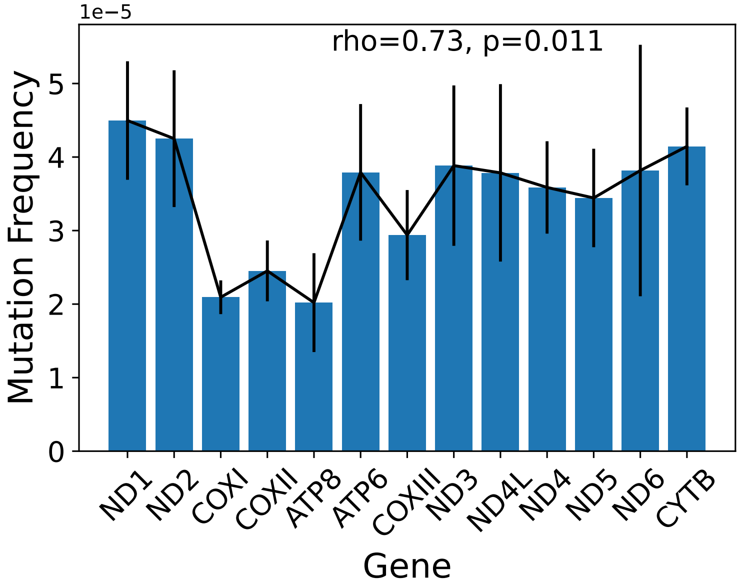


**Supplemental Figure 1. Per gene analysis of G→A mutation frequencies in aged (26mo) kidney sorted by gene position.** Mutation frequency was calculated by dividing the number of G→A mutations within the specified gene by the number of genes sequenced in the same gene. Error bars denote standard deviation. Correlation determined by Spearman's correlation.

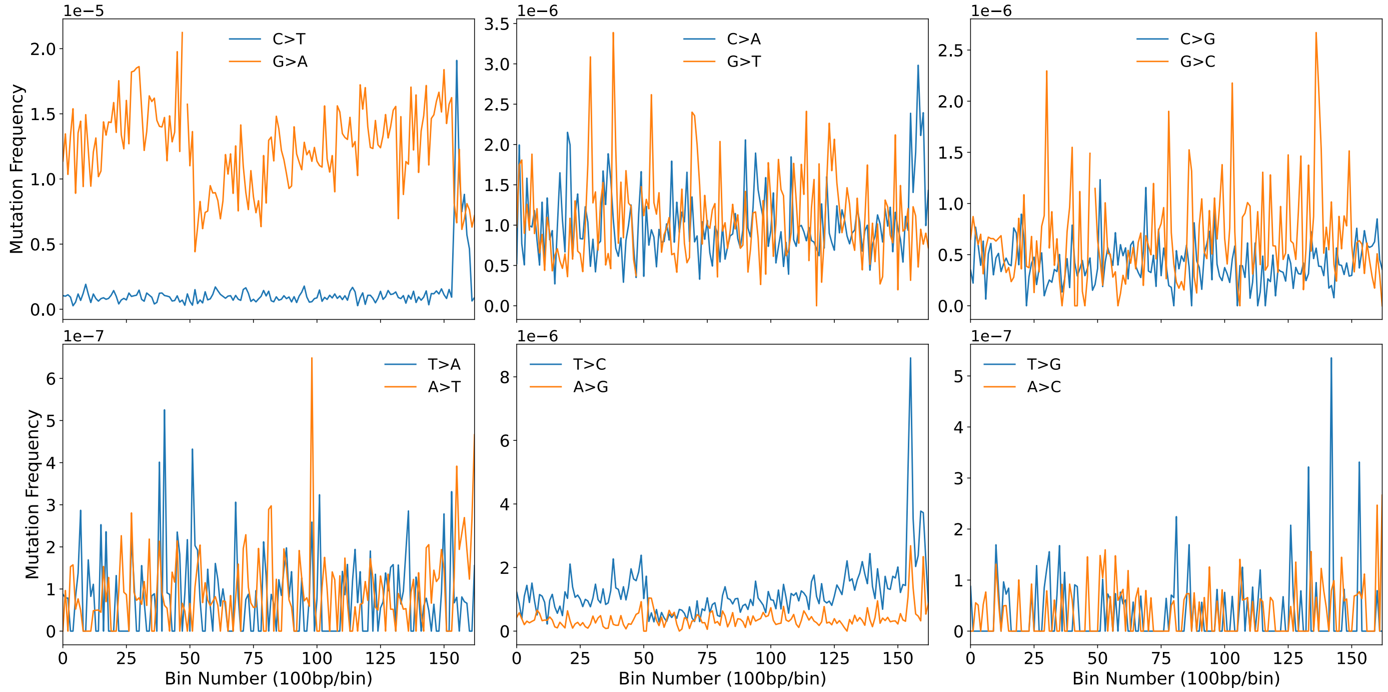


**Supplemental Figure 2. Distribution of all 12 mutation classes across the wild-type mouse mtDNA.** Mutations are reported as found on the L-strand. Complementary mutation types are grouped together in the same pane. Order of magnitude for the bin-specific mutation frequency for reciprocal types is located at the upper left of each pane.

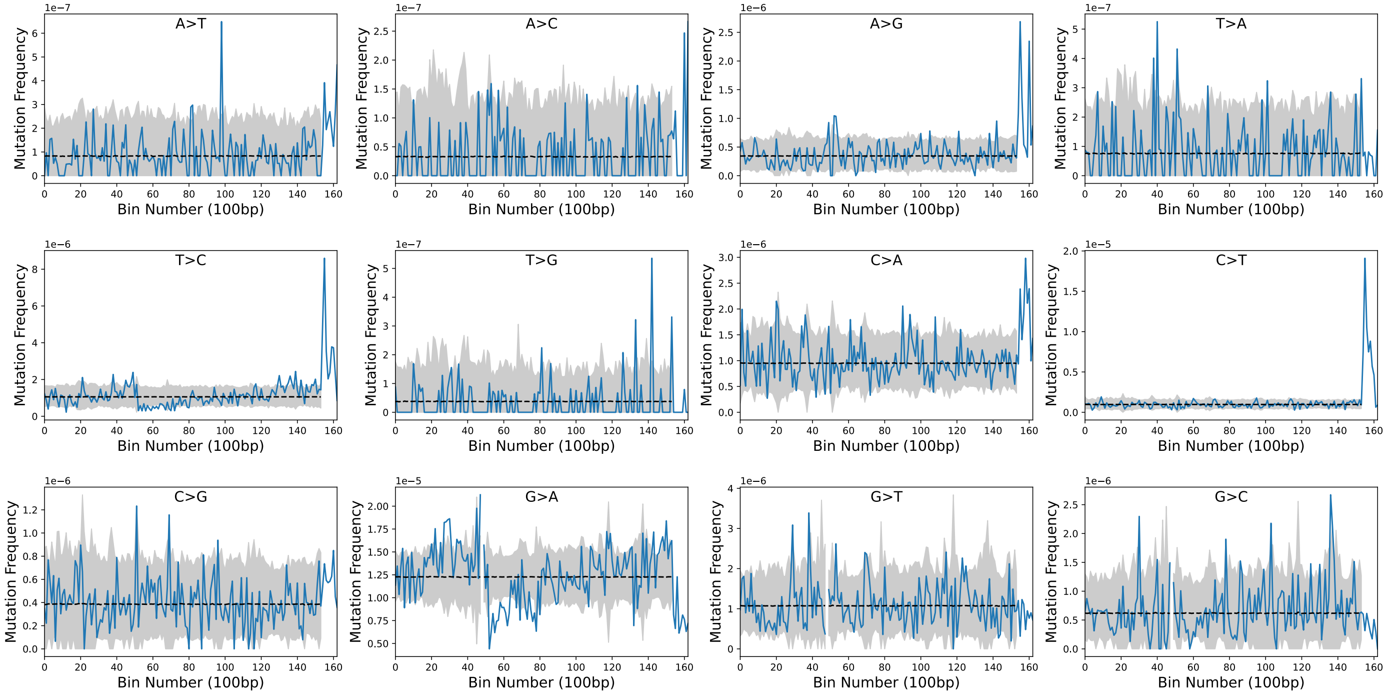


**Supplemental Figure 3. Simulated somatic mutation frequencies across mouse mtDNA for each mutation class.** The observed bin-specific mutation frequencies is in blue and are the same as reported in Supplemental Figure 1. The simulated bin-specific mean is reported as a dashed line. The 95% confidence interval of the 10,000 simulations is shaded in gray. Order of magnitude for each pane is located in the upper left corner.

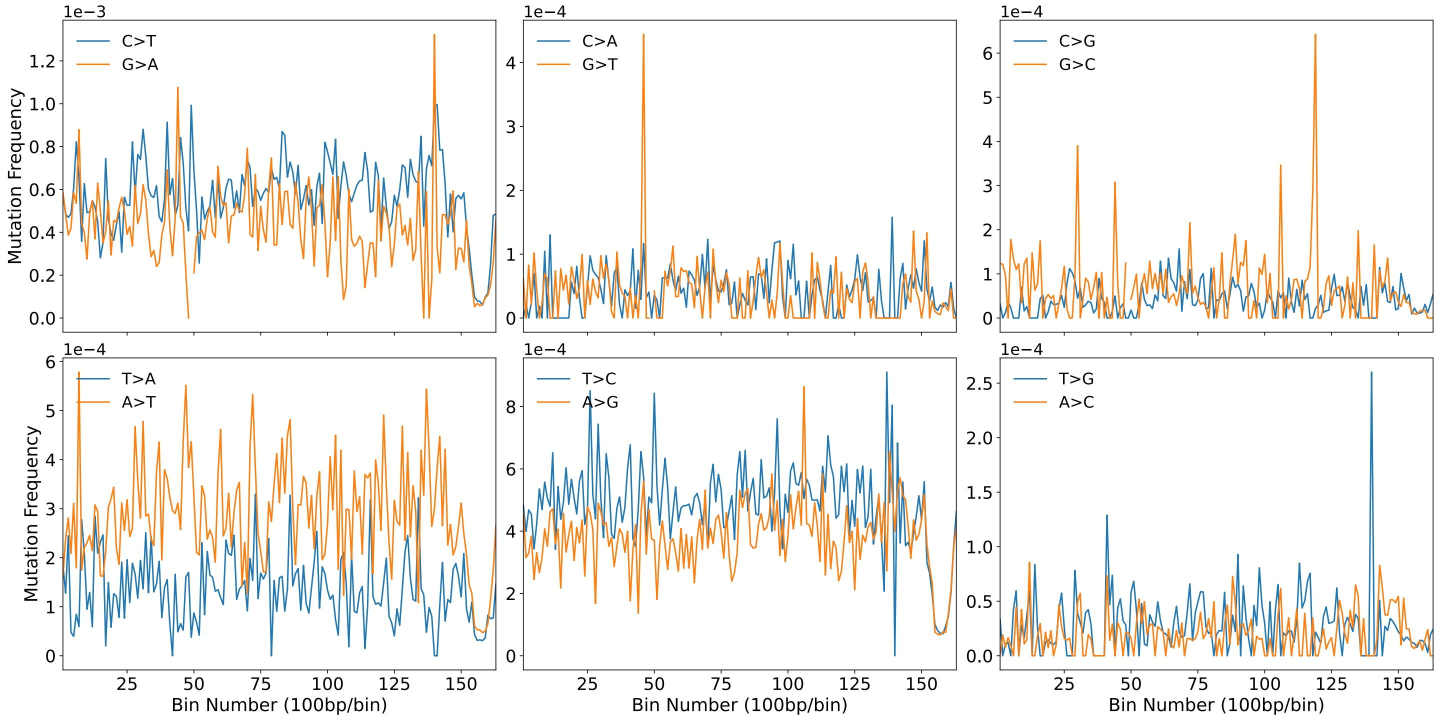


**Supplemental Figure 4. Distribution of all 12 mutation classes across the Pol-γ$^{exo-}$ mouse mtDNA.** Mutations are reported as found on the L-strand. Complementary mutation types are grouped together in the same pane. Order of magnitude for the bin-specific mutation frequency for reciprocal types is located at the upper left of each pane.

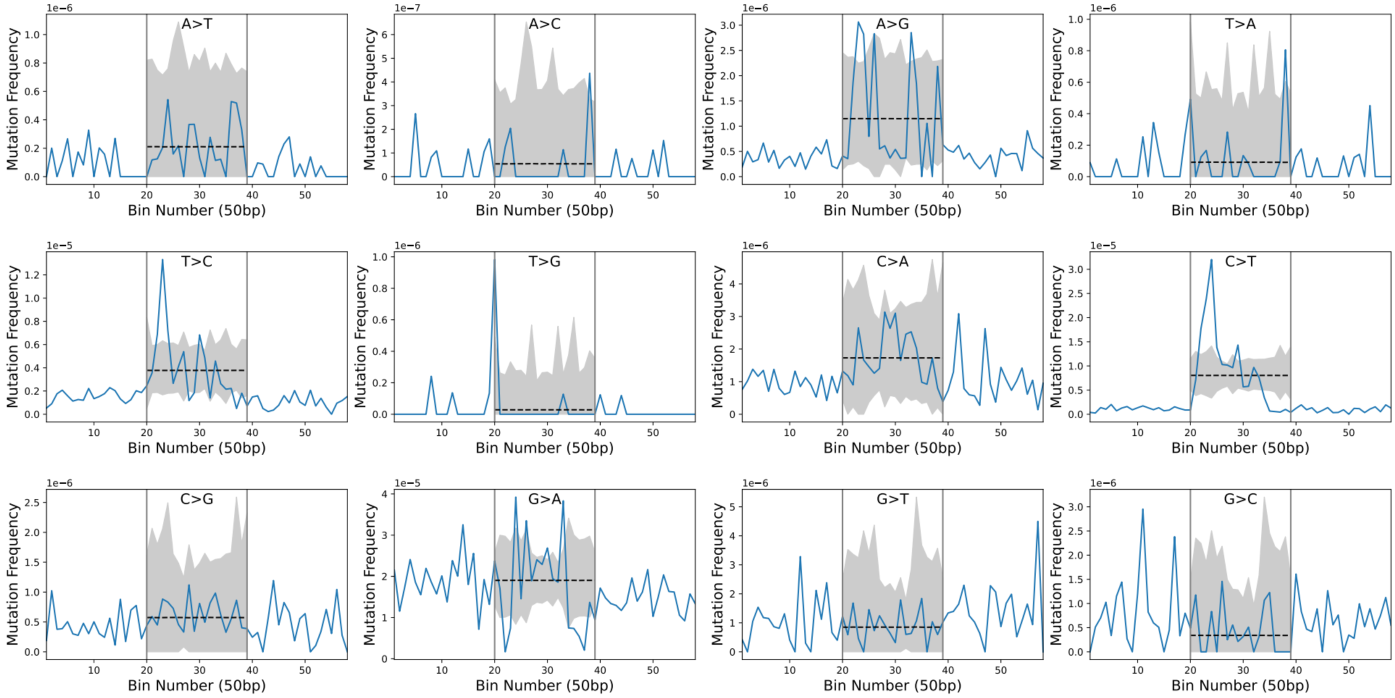


**Supplemental Figure 5. Simulated distribution of mutations in the mouse mtDNA control region.** The observed bin-specific mutation frequencies is reported as the blue line. The simulated bin-specific mean is reported as a dashed line. The corrected confidence interval (99.975%) of 100,000 simulations is shaded in gray. Any observed bin-specific mutation frequencies outside of the shaded region are over- or under-represented. Order of magnitude for each pane is located in the upper left corner. Vertical gray lines denote boundaries of the bins used for the simulation and are the nearest bin encompassing any sequence of the CR.

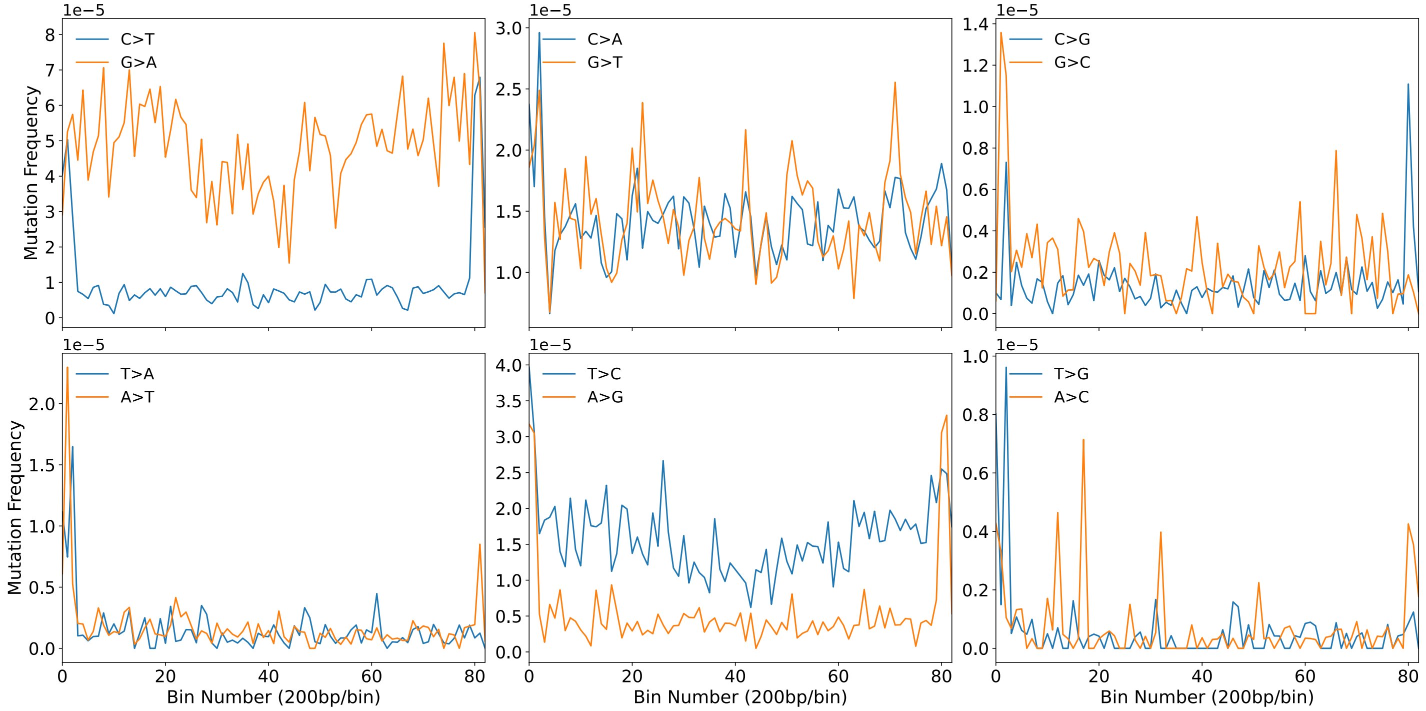


**Supplemental Figure 6. Distribution of all 12 mutation classes across the human mtDNA.** Mutations are reported as found on the L-strand. Complementary mutation types are grouped together in the same pane. Order of magnitude for the bin-specific mutation frequency for reciprocal types is located at the upper left of each pane.

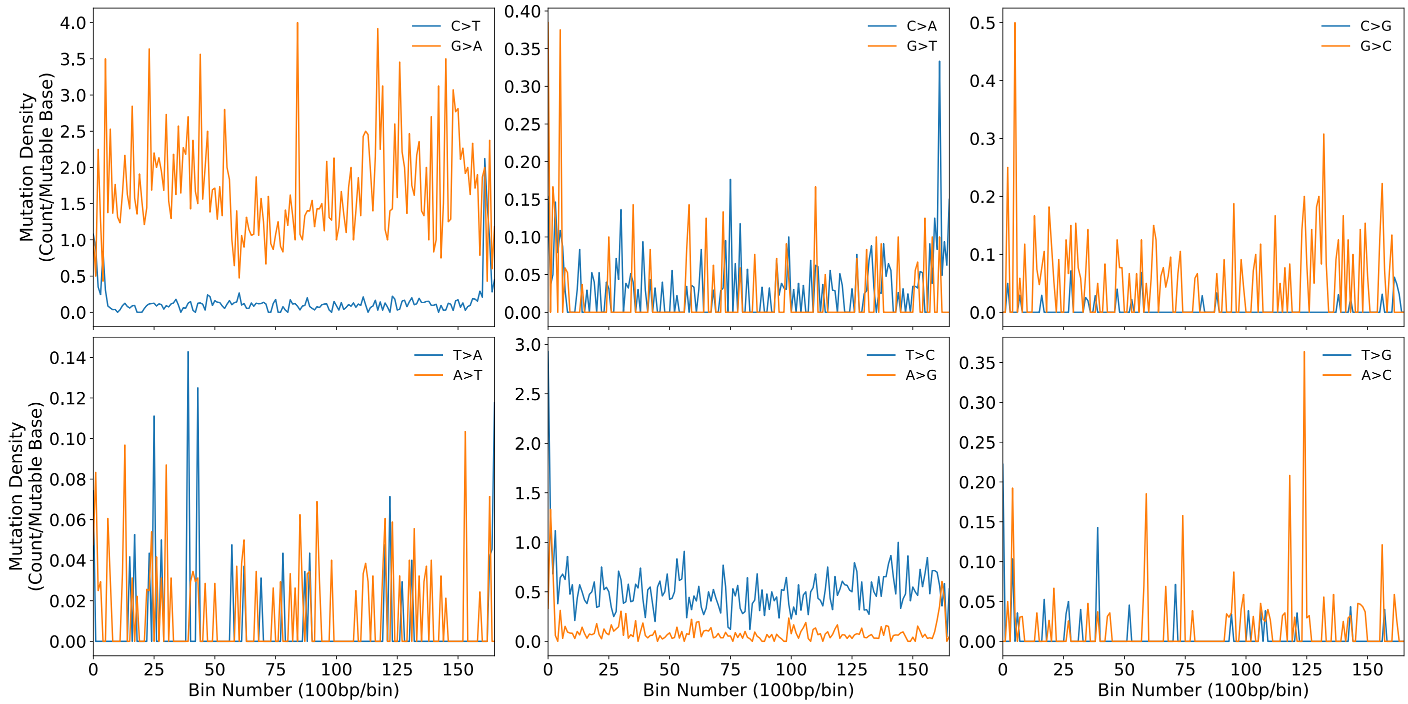


**Supplemental Figure 7. Distribution of all 12 mutation classes across the human mtDNA as reported in the PCAWG data.** Mutations are reported as found on the L-strand. Complementary mutation types are grouped together in the same pane.

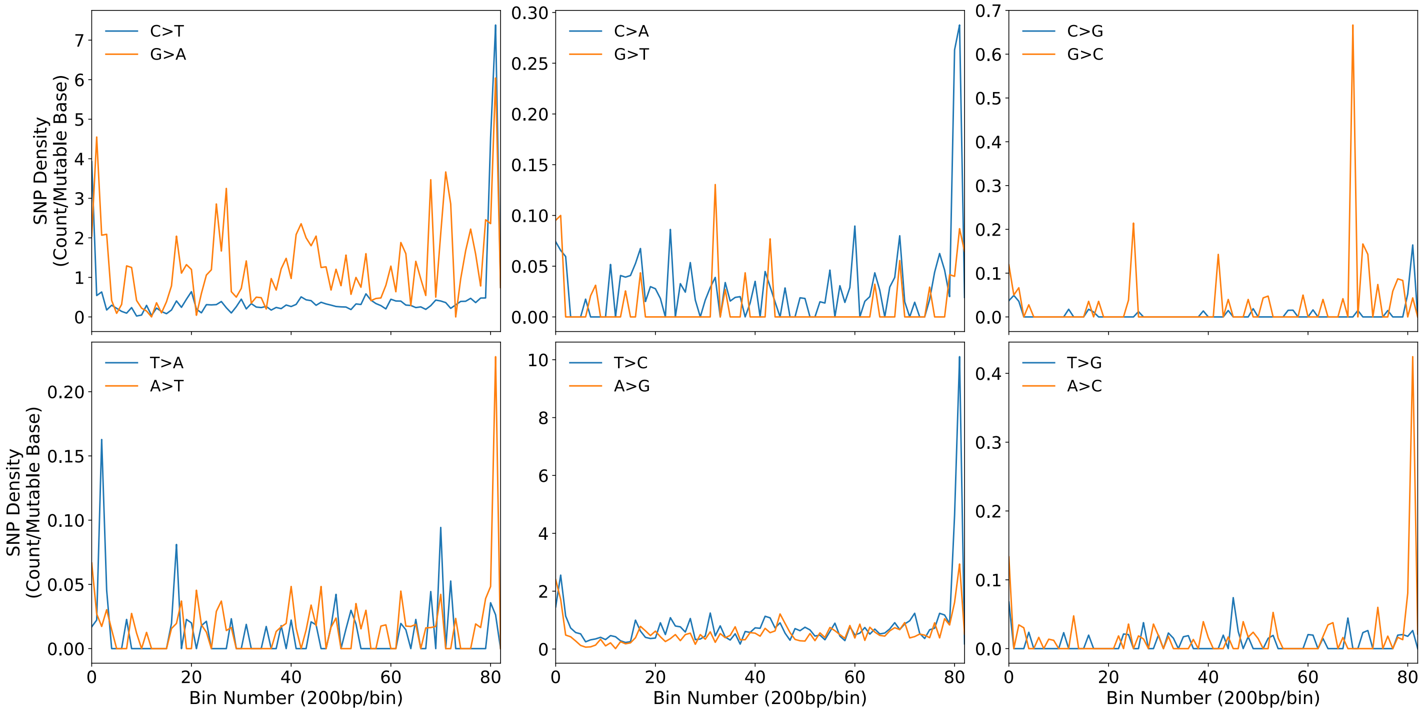


**Supplemental Figure 8. Distribution of SNPs from all 12 mutation classes across the human mtDNA.** Mutations are reported as found on the L-strand. Complementary mutation types are grouped together in the same pane. Data are taken direct from Supplemental Table 2 in Gu *et al.* [40].

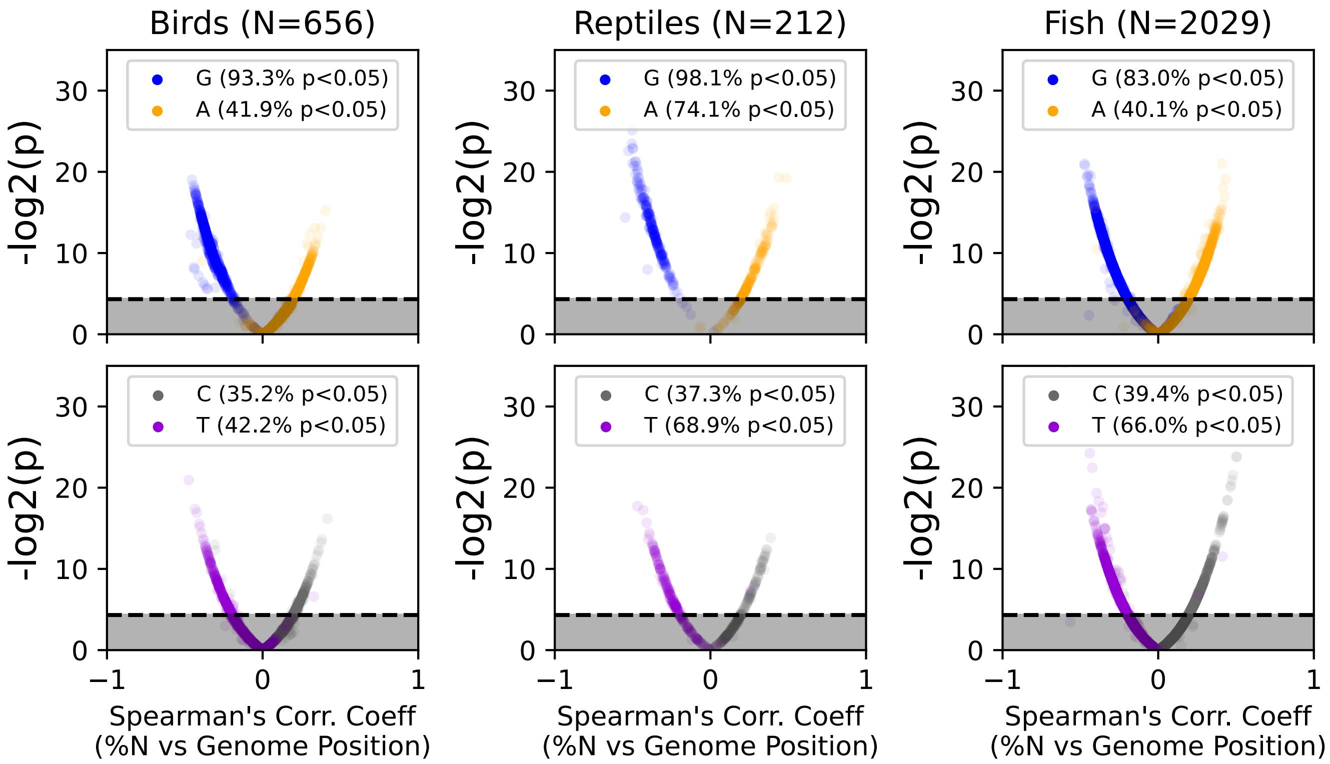


**Supplemental Figure 9. Volcano plot significant correlation between base composition and bin number across mtDNA sequences of diverse vertebrate species.** Base correlations are vertically grouped by taxonomic Class. Gray shaded area denotes non-significant correlation (p>0.05). Dotted line denotes p=0.05.

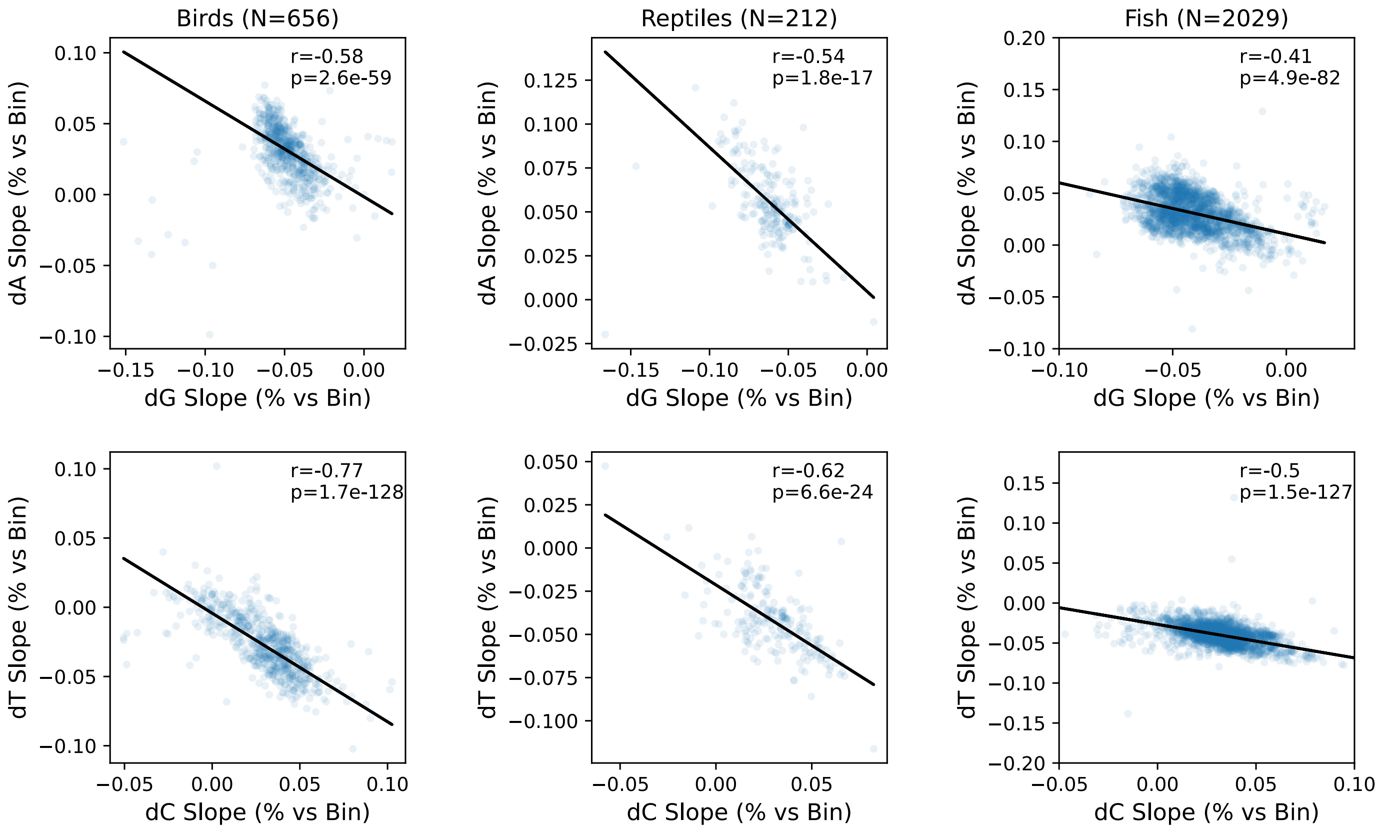


**Supplemental Figure s10. Correlation of slopes base composition vs number across mtDNA sequences of diverse vertebrate species.** Base correlations were determined by Spearman's correlation and grouped vertically by taxonomic Class with dG vs dA on the top and dT vs dC on the bottom. Black line denotes best fit by a robust linear model as described in the methods.
